# Minimizer Density revisited: Models and Multiminimizers

**DOI:** 10.1101/2025.11.21.689688

**Authors:** Florian Ingels, Lucas Robidou, Igor Martayan, Camille Marchet, Antoine Limasset

## Abstract

High-throughput sequence analysis commonly relies on *k*-mers (words of fixed length *k*) to remain tractable at modern scales. These *k*-mer-based pipelines can employ a sampling step, which in turn allows grouping consecutive *k*-mers into larger strings to improve data locality. Although other sampling strategies exist, local schemes have become standard: such schemes map each *k*-mer to the position of one of its characters. A key performance measure of these schemes is their density, defined as the expected fraction of selected positions. The most widely used local scheme is the minimizer scheme: given an integer *m* ≤ *k*, a minimizer scheme associates each *k*-mer to the starting position of one of its *m*-mers, called its *minimizer*. Being a local scheme, the minimizer scheme guarantees covering all *k*-mers of a sequence, with a maximal distance between selected positions of *w* = *k* − *m* + 1. Recent works have established near-tight lower bounds on achievable density under standard assumptions for local schemes, and state-of-the-art schemes now operate close to these limits, suggesting that further improvements under the classical notion of density will face diminishing returns. Hence, in this work, we aim to revisit the notion of density and broaden its scope.

As a first contribution, we draw a link between density and the distance between consecutive selected positions. We propose a probabilistic model allowing us to establish that the density of a local scheme is exactly the inverse of the expected distance between the positions it selects, under the minimal and only assumption that said distances are somehow equally distributed. We emphasize here that our model makes no assumptions about how positions are selected, unlike the classical models in the literature. Our result introduces a novel method for computing the density of a local scheme, extending beyond classical settings.

Based on this analysis, we introduce a novel technique, named multiminimizers, by associating each *k*-mer with a bounded set of candidate minimizers rather than a single one. The candidate furthest away (in a precise sense defined in the article) is selected. Since the decision is made by taking advantage of a context beyond a single *k*-mer, this technique is not a local scheme — and belong to a novel category of meta schemes. Using the multiminimizer trick on a local scheme reduces its density at the expense of a controlled increase in computation time. We show that this method, when applied to random (hash-based) minimizers and to open-closed mod-minimizers, approaches a density of 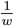, representing, to our knowledge, the first construction converging to this limit.

Our third contribution is the introduction of the *deduplicated density*, which measures the fraction of distinct minimizers used to cover all *k*-mers of a set of sequences. While this problem has gained traction in applications such as assembly, filtering, and pattern matching, standard minimizer schemes are often used as a proxy, blurring the distinction between the two objectives (minimizing the number of selected positions or the number of selected minimizers). Although related to the classical notion of density, deduplicated density differs in both definition and suitable constructions, and must be analyzed in its own right, together with its precise connections to standard density. We show that multiminimizers can also improve this metric, but that globally minimizing deduplicated density in this setting is NP-complete, and we instead propose a local heuristic with strong empirical behavior.

Finally, we show that multiminimizers can be computed efficiently, and provide a SIMD-accelerated Rust implementation together with proofs of concept demonstrating reduced memory footprints on core sequence-analysis tasks. We conclude with open theoretical and practical questions that remain to be addressed in the area of density.

## 1 Introduction

Efficient sequence comparison is a central computational pillar of modern genomics. Modern applications must process genomes of hundreds of gigabases, handle terabase-scale sequencing datasets, and search petabase-scale reference collections with both speed and tight memory usage. To perform sequence alignment at large scales, heuristic strategies such as seed-and-extend and chaining [1,2] avoid full dynamic-programming alignment, and prefer dividing this procedure into smaller or simpler tasks for which exact *k*-mer (a word of fixed length *k*) matches serve as high-confidence landmarks that guide the alignment. A key observation is that, in practice, one does not need to index all *k*-mers to maintain high alignment sensitivity; therefore, *k*-mer sampling is often preferred to decrease resource usage [3]. However, naive sampling schemes lack robustness to mutations or noise, and sampling schemes designed for distance approximation can generate “deserts” where no *k*-mer is chosen, leaving the alignment algorithm without the necessary anchor points [4]. Minimizers were introduced in this context [5,6] to couple sparsity (through sampling) with coverage guarantees (to avoid deserts as much as possible). This paper addresses several fundamental aspects of minimizers and their effectiveness in sequence representation.

Given a window of *w* = *k* − *m* + 1 consecutive *m*-mers (*k* ≥ *m*), a minimizer scheme selects one *m*-mer in this window according to an order, typically induced by a hash function. This ensures that any exact match of length at least *k* = *w* + *m* − 1 between two sequences yields at least one common minimizer, provided they use the same ordering. At first glance, this guarantees coverage, but not efficient sampling, since in principle each overlapping window could select a different *m*-mer. In practice, successive windows share *w* − 1 *m*-mers and differ by only one entering and one leaving *m*-mer, so they tend to select the same minimizer over long stretches. To formalize this, the notion of density was introduced: the average number of selected *m*-mers per window, or equivalently, the fraction of *m*-mers retained. This paper focuses on minimizer density, in particular in its optimization. To be explicit, a lower density means that fewer *m*-mers are selected, which directly reduces memory usage and comparison cost, or expectedly more cache-coherent algorithms, making it both a theoretically interesting question and one with broad practical impact, described succinctly below.

A wide range of tools now rely on minimizers as their main alignment and indexing mechanism. Long-read [7,8] and whole-genome aligners [9] index only minimizers for their heuristics, reducing memory footprint and accelerating chaining. Classifiers and document-retrieval indexes use minimizers to shrink database size by orders of magnitude [10] or to speed up filtering [11,12]. Lossy representations encoding sequences as ordered lists of minimizers have also become central in lightweight assembly and comparison methods [13,14]. Beyond window-based selection, minimizer schemes can assign to each *k*-mer one of its *m*-mer (*m* ≤ *k*) as its minimizer, inducing a natural partition of the *k*-mers without additional metadata. Consecutive *k*-mers often share the same minimizer, forming long runs that can be stored compactly as a single superstring of length *k* + *m* − 1 [15]. This property is exploited in *k*-mer counters [16,17], scalable de Bruijn graph construction [18,19,20], and large-scale indexing structures [21,22]. In this setup, the density matters again: higher density breaks these runs more often, reducing the benefits of the compact string representation.

Minimizers are typically computed on a stream of genomic sequences. A first key notion is that of local schemes, in which *m*-mers are selected solely based on the content of each window. A second notion is that of forward schemes, which restrict selection to *m*-mers appearing at increasing positions along the sequence. Every forward scheme is therefore a local scheme. With this window structure in place, the best possible density is 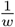 (one selected *k*-mer per window). Early work established random-order minimizers, e.g. hashing-based orderings, as a simple baseline with expected density 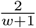, and conjectured that this might be near-optimal for local schemes [5]. Subsequent results disproved this, and multiple research threads have since pushed densities closer to the theoretical limit:

– Marçais et al. characterized when asymptotically optimal minimizers exist and introduced a practical construction [23].
– Another line of work proposed precomputing a set of minimizers that hits every possible window in the universe; so that a given input sequence then only selects minimizers from this set. These schemes improve density but suffer from exponential growth in construction cost as *k* increases. Pellow et al. addressed this with heuristics that scale to arbitrarily large *k* and any window length [24].
– Groot Koerkamp and Pibiri [25,26] introduced schemes based on modular constraints on internal subwords, achieving asymptotically optimal density 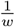 as *k* → ∞.

Kille and Groot Koerkamp et al. later derived a nearly tight lower bound for forward sampling schemes, showing that for many practical (*k, w*), state-of-the-art constructions are within only a few percent of optimal density [27]. This suggests that substantial further improvements in local schemes are unlikely. Recent schemes such as Golan et al. [28] attain densities extremely close to the Kille and Groot Koerkamp et al. bound for practical parameter ranges, as do the aforementioned constructions of Pellow et al., and Groot Koerkamp and Pibiri.

To escape the limitations of universal forward local schemes, which are designed to perform well on any input, the community has also designed sequence-specific schemes that tailor minimizer selection to the properties of a given genome. These include frequency-based selection [29], spacing-based strategies [30], and machine-learning approaches [31]. While these methods can substantially reduce density on real genomes, they cannot be computed in a streaming fashion, making them impractical at very large scales.

Despite the extensive effort devoted to low-density minimizer schemes, several aspects of density behavior remain poorly understood. In this work, we focus on three such aspects and use them as the foundation for our contributions.

First, we revisit classical density for local schemes in a unified way. We formalize the link between density and distances between consecutive selected positions and show that, under a minimal assumption on these distances, the density of *any* local scheme equals the inverse of the expected distance. Specializing to random minimizers, we argue that the standard modeling assumption is unsatisfactory. We thus propose a more realistic one that coincides with the classical regime for large *m*-mers, and verify empirically that it satisfies our minimal condition.

Second, we build on this perspective to design practical family of meta schemes dubbed *multiminimizers*. Instead of fixing a single minimizer per *k*-mer, we assign a bounded set of candidate minimizers and choose among them to reduce the frequency of selected positions while preserving locality and consistency. Instantiated on top of a random-hash order, multiminimizers attain asymptotically optimal positional density, match or surpass the best known schemes in practice, and remain simple and efficient to implement.

Third, we make explicit a twin notion to classical density: the *deduplicated density*, the fraction of distinct minimizers needed to cover all *k*-mers, directly relevant for filters and indexes. We show that classical and deduplicated densities diverge on long random sequences, but essentially coincide for small sequences, when computed for random minimizers. We then study minimizing deduplicated density in the multiminimizer framework, prove that the global optimization problem is NP-complete (via reduction from set cover), and introduce a local heuristic with strong empirical behavior. This detour clarifies the relationship between the two densities and highlights a remaining open problem: establishing the computational complexity of minimizing classical density itself.

## 2 Methods

### 2.1 Preliminaries

In this section, we introduce key concepts for the rest of this article. We build mainly on the notations of [17,32,25,33,28]. We are interested in strings defined over the DNA alphabet `{A, C, G, T}` of size *σ* = 4. Let *S* ∈ *Σ*^∗^ be a string; we denote by *S*[*i* … *j*) the substring of *S* of length *j* − *i*, starting at position *i* (included) and ending at position *j* (excluded). A *k*-mer is a substring *S*[*i* … *i* + *k*) of fixed length *k*. The length of a string *S* is denoted by |*S*| . Most of the time, the DNA strand from which a sequence is coming is unknown; the ambiguity is resolved by considering that a *k*-mer and its reverse complement are, in fact, the same *k*-mer. One of the two (e.g. the lexicographically smallest) is chosen to be the *canonical* representative of the pair. Hereafter, the definitions are stated in a non-canonical context (but are easily extended); whereas the practical experiments presented later on are based on canonical *k*-mers, although our implementations cover both contexts. On the topic of canonical *k*-mers, see also [4].

An order 𝒪_*m*_ on *m*-mers is an injective function *Σ*^*m*^ → ℝ such that 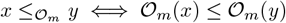. In practice, we define an order using a (pseudo)-random hash function *h* : *Σ*^*m*^ → *U* [34], where *U* is large enough so that the probability of collision (i.e. *h*(*x*) = *h*(*y*) with *x* ≠ *y*) can be assumed to be 0 – i.e. *h* is a perfect hash function. Typically, *U* = 2^64^ whenever using 64-bits hash functions. We denote by 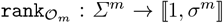 the function that associates to a *m*-mer its rank among all *m*-mers according to the order 𝒪_*m*_. Since 𝒪_*m*_ is injective, 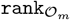 is a bijection.

#### Definition 1

**((Random) Minimizer)**. *The minimizer* min(*x*) *of a k-mer x is its smallest substring of size m < k for the (random) order* 𝒪_*m*_.

Let us denote *w* = *k* − *m* + 1 the number of *m*-mers contained in a *k*-mer.

#### Definition 2

**(Minimizer scheme)**. *Provided an order* 𝒪_*m*_, *a minimizer scheme is a function f* : *Σ*^*k*^ → ⟦1, *w* ⟧ *that returns the position of the minimizer of a k-mer, with ties resolved to the left; i*.*e. f* (*x*) = min{1 ≤ *i* ≤ *w* : *x*[*i* … *i* + *m*) = min(*x*)}.

#### Definition 3

**(Set of selected indices)**. *The set of selected indices of a string S using the minimizer scheme f is defined as* 𝒮_*f*_ (*S*) = {*i* + *f* (*S*[*i* … *i* + *k*)) : 1 ≤ *i* ≤ |*S*| − *k* + 1}.

As noted in [33], the scheme may select the same position in consecutive windows (since consecutive windows overlap by *k* − 1 bases), so that |𝒮_*f*_ (*S*)| ≤ |*S*| − *k* + 1.

#### Definition 4

**(Density, [28])**. *The (expected) density of a minimizer scheme f is formally defined as*

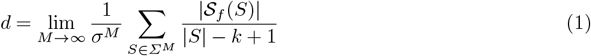

*and represents the expected proportion of selected minimizers among all m-mers on a random string, where each character is chosen uniformly at random. For any sequence S, the term* 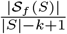 *is also defined as the particular density of S, and denoted by d*(*S*) . *Since* 1*/w* ≤ *d*(*S*) ≤ 1, *we also have* 1*/w* ≤ *d* ≤ 1.

Marçais et al. proved that the density can be computed exactly (instead of just being estimated) by noticing that the density and the particular density coincided when computed on a de Bruijn sequence of large enough order [35]. The purpose of the upcoming Section 2.2 is to provide yet another way to compute the density. Finally, note that all these definitions can also be applied to the broader concept of *local schemes*, which are guaranteed to select one *m*-mer per *k*-mer. Minimizer schemes are one example of local schemes, see also [33].

### 2.2 A link between density and the expected distance between selected positions

Let *f* be a minimizer scheme. Consider the expected number of selected positions in a window of size *w* — by definition, is is equal to *d* · *w*. Intuitively, this quantity is also equal to *w* divided by the expected distance between two selected consecutive positions — that we denote here by *µ*. Indeed, if we select a position every *µ* bases, we expect to choose *w/µ* positions on average in a window. Hence, with *dw* ≈ *w/µ*, we expect 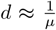. This intuitive link has already been noticed in the literature [5,30,17], but never, to the best of our knowledge, explicitly formalized as such, nor proved for any scheme. The purpose of this section is to formally establish this observation as a theorem, namely Theorem 1, under minimal assumptions.

The sequence *S* is represented by a vector of integers **R** = *R*_1_, *R*_2_, …, where 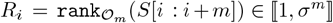 is the rank of the *i*-th *m*-mer of *S*, according to the order 𝒪_*m*_. A second integer vector, **P** = *P*_1_, *P*_2_, …, describes the positions of the selected minimizers when sliding a window of size *w*, i.e.

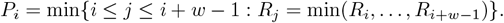

Note that **P** can still be defined no matter how minimizers are chosen, or more generally, a position, in a window. Since consecutive windows may choose the same position as a minimizer, consecutive values of *P*_*i*_ might be equal. It leads us to define the subsequence **P**^∗^ of *distinct* values, i.e. 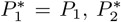 is the first *P*_*i*_ value distinct from 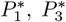 is the first *P*_*i*_ value distinct from 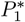 and 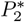, and so on. In other words, **P**^∗^ is the sequence corresponding to the set 𝒮_*f*_ (*S*) – i.e. the set of selected indices. Finally, we define **Δ** = *Δ*_1_, *Δ*_2_, … as the sequence of distances between selected positions, i.e. 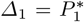, and 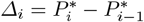. An example is provided in Figure 1.

**Fig. 1:**
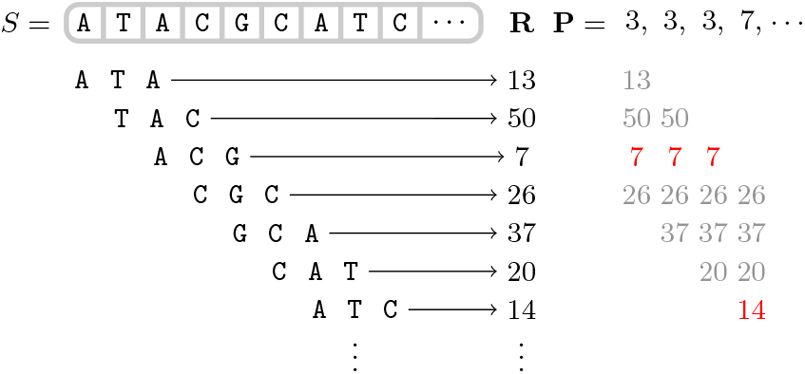
An example of sequence *S*, represented as a sequence **R** of integers (here, using lexico-graphical order), and the sequence **P** of minimizers positions, with *m* = 3 and *w* = 4. We have **P**^∗^ = 3, 7, … and **Δ** = 3, 4, … .

We put ourselves in the context where *S* is a random sequence, so that **R, P, P**^∗^ and **Δ** can be considered as vectors of random variables. Our goal is to establish a link between *d* and the expected distance between selected positions, i.e., between *d* and the 𝔼[*Δ*_*i*_]’s. We obtain the following result.

#### Theorem 1.

*Let µ >* 0. *Consider ε*_*i*_ = 𝔼[*Δ*_*i*_] − *µ as some realization of an underlying random variable ε with* 𝔼 [*ε*] = 0. *Then, we have d* · *µ* = 1.

*Proof*. The proof is deferred to Appendix S1.

Basically, Theorem 1 states that if we assume that the distances between consecutive selected positions of minimizers are somehow equally distributed, then their expected value is exactly 1*/d*. This assumption is the minimum viable hypothesis for establishing the result (since *µ* must be well defined), and note also that this hypothesis does not depend on the process by which the *R*_*i*_’s values are obtained, nor how the positions of the minimizers *P*_1_, *P*_2_, … are selected in successive windows — thus making our result very general, applicable to *any local scheme*, and, to the best of our knowledge, the first of its kind.

At this point, we have not made any assumptions on the random vectors **R, P** and **P**^∗^. It is not trivial to formally verify whether the assumption regarding the 𝔼[*Δ*_*i*_]’s is verified or not, as the variables are most likely not to be independent nor identically distributed. However, in practice, we can verify that the assumption holds. In the remainder of this section, we investigate how the assumption from Theorem 1 concerning the *Δ*_*i*_’s holds for random minimizers. The standard hypothesis that has been made in the literature regarding their choice is the following:

#### Hypothesis 1

**([5,6,35])** *Every m-mer in a window of length w has an equal probability of* 1*/w of being the smallest m-mer*.

As discussed in [35], while “not strictly true in practice”, this hypothesis is reasonable and reflects reality accurately when using a randomized ordering”. However, the formulation is unfortunate because it does not take into account the dependency between windows. If we consider a window (*R*_*i*_, …, *R*_*i*+*w*−1_) outside of its context, then the hypothesis can be reformulated as *P*_*i*_ ∼ 𝒰 (⟦1, *w*⟧). Unfortunately, since the same minimizer will likely be selected by several successive windows, *P*_*i*_ cannot be considered independent, nor even identically distributed (since *P*_*i*_ = *P*_*i*−1_ with a non-zero probability). Here, we propose starting from a more elementary hypothesis concerning the *R*_*i*_’s and observing, empirically and theoretically, the behavior of the variables derived from them, **P, P**^∗^ and **Δ**, in order to study the difference between Hypothesis 1 and reality, as well as the hypothesis concerning the expectation of the *Δ*_*i*_’s. Our working assumption is the following.

#### Hypothesis 2

*The random variables R*_1_, *R*_2_, … *are i*.*i*.*d, uniformly distributed over* ⟦1, *σ*^*m*^ ⟧.

*Remark 1*. Assuming a perfect hash function when computing the random order, our assumption states that the probability of collision between *R*_*i*_’s is 1*/σ*^*m*^ — which is consistent with [30, Lemma S7], that establishes that two distinct *m*-mers are identical in a random sequence with probability 1*/σ*^*m*^.

It is not too difficult to establish the law of 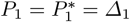 from Hypothesis 2:

#### Proposition 1.

*Under Hypothesis 2*, ∀1 ≤ *i* ≤ *w*,

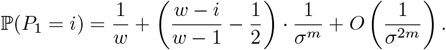

*It follows that* 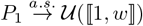 *when m* → ∞.

*Proof*. The proof is deferred to Appendix S2.

In Appendix S3, we observe that the limit distribution is reached very quickly, for fairly small values of *m* (*m* = 8 in our case), which implies that for all values of *m* likely to be used in practice, Hypothesis 1 is verified for *P*_1_, and *Δ*_1_ also follows a uniform distribution, meaning that 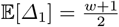. If, indeed, all subsequent *Δ*_*i*_’s have the same expectation, then, using Theorem 1, we retrieve the well-known density of random minimizers, 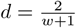 [5,6].

Unfortunately, formally establishing the law of further variables becomes rapidly intractable due to the dependencies between them. To complete the study of subsequent 𝔼[*Δ*_*i*_]’s, we resort to Monte-Carlo simulations of our probabilistic model and obtain the results depicted in Appendix S4. From our results, the assumption that 𝔼[*ε*] = 0 is empirically verified (numerically, we get 𝔼 *ε*] ≈ 0.0060), and the 𝔼 [*Δ*_*i*_]’s are indeed distributed around 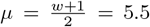 (using *w* = 10) — see supplementary Figure 8a. We obtain numerically a density of 0.1808, i.e. 0.55% of error compared to the theoretical value of 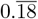. This empirical result highlights that Hypothesis 2 is indeed a proper assumption to work with, and also that the assumption of Theorem 1 is reasonable, since it is verified in practice.

### 2.3 Multiminimizers: trading time for space

Considering Theorem 1, obtaining a low density scheme on a sequence *S* requires having a scheme that selects positions as far as possible from one another in *S*, while still covering all *k*-mers from *S*. In this regard, the best possible minimizer scheme would select *m*-mers at a distance of *k* − *m* + 1 bases from each other. However, without a way to retrieve which *m*-mer was chosen for a specific *k*-mer, querying a *k*-mer would require checking if any of its *k* − *m* + 1 *m*-mers was selected as its minimizer. The same problem arises when inserting a *k*-mer into a database: one must perform *k* − *m* + 1 checks to verify whether it is already present or not.

To reduce this prohibitive cost, we propose a simple yet efficient solution leveraging the notion of super-*k*-mers. A *super-k-mer* is defined as a maximal sequence of consecutive *k*-mers having the same minimizer [36]. Our technique starts by generating *N* distinct hash functions (e.g. using *N* distinct random seeds). Then, we use these *N* hash functions to generate *N* distinct random minimizer schemes. Each of these minimizer schemes yields a set of super-*k*-mers on the sequence *S*, each covering all *k*-mers of *S*. When iterating over the minimizers of a sequence, among the super-*k*-mers that cover the first *k*-mer, we select the one that ends the farthest in the sequence. Then, we repeat this selection at the end of each selected super-*k*-mer: among all the super-*k*-mers covering the first uncovered *k*-mer, we select the one that ends the farthest in the sequence (see Figure 2). We detail our method in Algorithm 1, running in *O*(*N* · |*S*|) time and using *O*(*N* · *w*) memory.

**Fig. 2:**
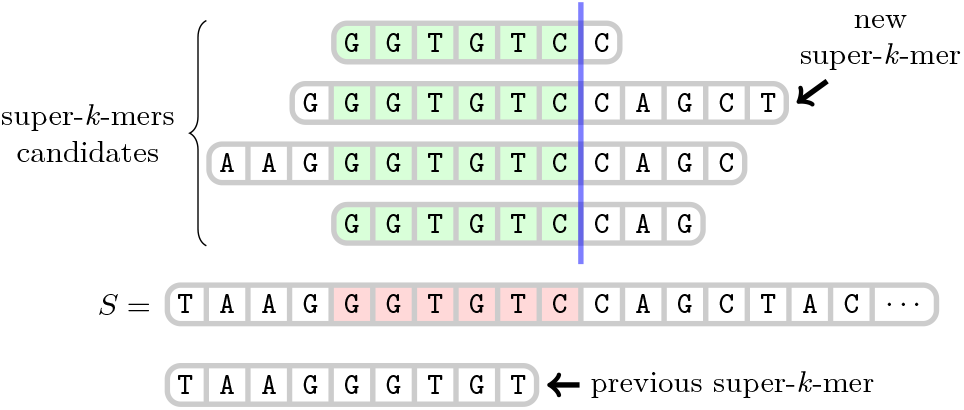
Choice of a super-*k*-mer for *k* = 6 using a multiminimizer scheme. The first *k*-mer uncovered by the previous super-*k*-mer is highlighted in red. Among the *N* = 4 candidates super-*k*-mers covering this *k*-mer, the multiminimizer scheme selects the one ending the farthest in the sequence (i.e. farthest right of the blue line).

As mentioned in the introduction, a *local scheme* selects a *m*-mer in a *k*-mer solely based on the *k*-mer itself. This ensures covering all *k*-mers, or equivalently, all windows of *w* = *k* − *m* + 1 *m*-mers [33]. Formally, a local scheme is a function *f* : *Σ*^*k*^ → ⟦1, *w* ⟧. Importantly, the local scheme is both used at construction (e.g. when partitioning *k*-mers based on their minimizer) and query (e.g. to determine in which bucket a *k*-mer might be present). For a given *k*-mer *x* ∈ *Σ*^*k*^, the position *f* (*x*) is uniquely determined. While minimizers are local schemes, multiminimizers are not, as they differ drastically in their approach:

– At construction, we need to know when the previous super-*k*-mer ends, and where the candidates super-*k*-mers end, informations that a *k*-mer alone cannot provide. In this regard, the multiminimizer scheme has both to “remember the past” to determine when to select a new minimizer, and “delve into the future” to choose which minimizer candidate to keep.
– At query, for a given *k*-mer, we have to compute and lookup the *N* minimizers candidates to determine whether it has already been seen, since the broader context(s) in which it might have been seen are not known.

In the end, neither at construction nor query does the multiminimizer scheme behave like a local scheme. It is advantageous as it implies that its density is not subject to the lower bound of local schemes by Kille and Groot Koerkamp et al. [27] — although the 1*/w* bound obviously remains. As showed in upcoming Section 3.1, the multiminizer scheme does indeed beat this lower bound, and approach density of 1*/w* as *N* increases. This gain in density comes at the price of (bounded) increased query time, as per the discussion above.

Finally, note that the multiminimizer trick is not limited to random minimizer schemes, but could be used with any combination of *N* distinct local schemes (whether by varying parameters or using concurrent schemes) — in that sense, multiminimizers are a sort of *meta* scheme. To illustrate this point, in upcoming Section 3, we investigate two implementations of the multiminimizer trick,one with random minimizers, and the other with open-closed mod-minimizers [26].

#### Algorithm 1

MultiMinimizers

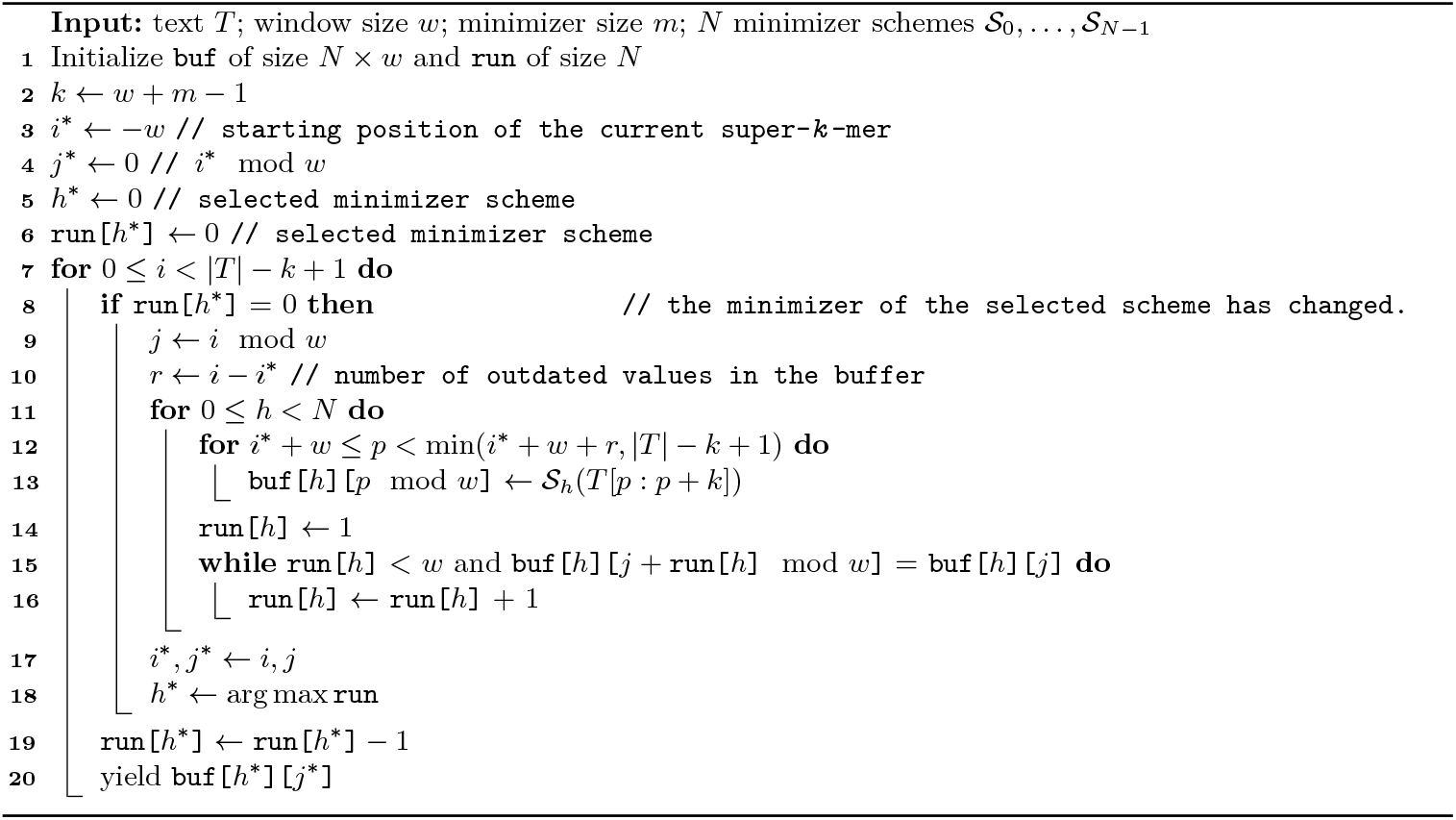

### 2.4 On deduplicated density

It appears the standard density possesses a “twin” concept, that we call *deduplicated density* (which we denote by *d*^∗^), and that we define as the fraction of distinct minimizers needed to cover all *k*-mers of a set of sequences (or, equivalently, a set of *k*-mers), instead of the fraction of positions selected along a sequence. To the best of our knowledge, the deduplicated density has never been mentioned in the literature, even if it is implicitly linked to filter problems, such as the ones tackled by Needle [37].

#### Definition 5

**(Minimizer Cover)**. *Let* 𝒳 = {*x*_1_, *x*_2_, … } *be a set of (distinct) k-mers. The* minimizer cover *of* 𝒳, *according to some minimizer scheme f, that we denote by M**in**C**over*_*f*_ (𝒳), *is defined as M**in**C**over*_*f*_ (𝒳) = {min_*f*_ (*x*) : *x* ∈ 𝒳}, *where* min_*f*_ (*x*) *designates the minimizer of x according to f* .

Analogously to standard density — recall Definition 4, we can define a *particular* deduplicated density *d*^∗^(𝒳) and obtain simple bounds, mimicking the ones we already have on standard particular density:

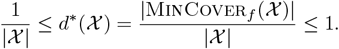

Standard density is computed as the limit of the expected particular density 𝔼 [*d*(*S*)] on a random sequence *S* when |*S*| → ∞, for fixed values of *k* and *m* — recall (1). However, one cannot have an arbitrarily large set 𝒳 since |𝒳 | ≤ *σ*^*k*^; therefore, we propose to define the deduplicated density simply as the expected particular density of a random set of *k*-mers 𝒳, without any notion of limit.

#### Definition 6.

*The (expected) deduplicated density of a minimizer scheme f is formally defined as*

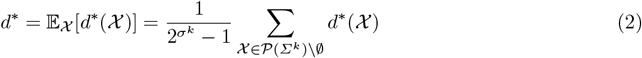

*where* 𝒫(*Σ*^*k*^) *designates the powerset of Σ*^*k*^.

Noticing that 1*/*|𝒳| ≥ 1*/σ*^*k*^, we obtain the following trivial bounds for *d*^∗^.

#### Lemma 1.

*For any minimizer scheme, we have* 1*/σ*^*k*^ ≤ *d*^∗^ ≤ 1.

Then, we have the following result.

#### Proposition 2.

*Under Hypothesis 2, the deduplicated density of the random minimizer scheme is given by the following, where w* = *k* − *m* + 1:

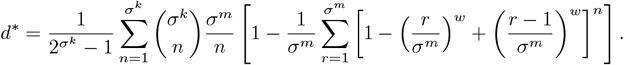

*Proof*. The proof is deferred to Appendix S5.

The dependence on *σ*^*k*^ is unfortunate, making it intractable to compute *d*^∗^ when *k* is large. We nonetheless computed the first values, shown in Figure 3 (left); one can see that *d*^∗^ seems to be driven by the value of *w* rather than *k*, hence we conjecture that *d*^∗^ might only depend on *w*. Besides, it seems that *d*^∗^ *< d* holds. To access higher values, we resorted to Monte-Carlo simulations, following the same probabilistic model as before, using Hypothesis 2. We observed how the standard density *d* and the deduplicated density *d*^∗^ evolved over the course of parsing a random sequence, as shown in Figure 3 (right). While both densities seem to be equal for the first few thousand windows, they rapidly diverge. We looked into the data and observed that the number of distinct *k*-mers and distinct windows observed are equal (it is not surprising since the probability of collision of *k*-mers is negligible), so the difference between the densities is entirely explained by the discrepancy between the number of selected positions and the number of distinct minimizers. While those values are equal at first, soon we obtain repeated minimizers^3^, hence the drop in *d*^∗^.

**Fig. 3:**
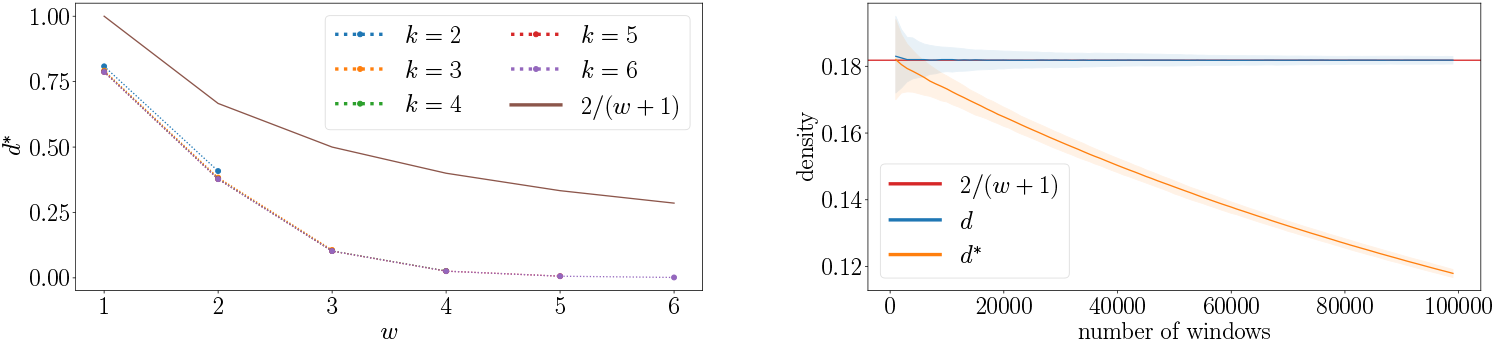
(left) Theoretical values of *d*^∗^ for 2 ≤ *k* ≤ 6 as a function of 1 ≤ *w* ≤ *k*, obtained with the formula of Proposition 2. (right) Monte-Carlo simulations of random minimizers under Hypothesis 2. We generated *N*_simu_ = 10^3^ random sequences **R** of size |**R**| = 10^5^, with *m* = 8 and *w* = 10. For *i* ≥ 1, we recorded the current values of *d* and *d*^∗^ estimated on the sequence every *i* 10^3^ windows. 95% of values across all simulations are within the colored areas, the solid lines being the medians.

There are obviously many more aspects of deduplicated density to be studied and discovered, but they are beyond the scope of this article. To conclude this section, we propose to consider applying multiminimizers to deduplicated density.

#### Definition 7

**(Multiminimizer Cover)**. *Let* 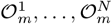 *be N orders on m-mers, and let* 𝒳= {*x*_1_, *x*_2_, … } *be a set of (distinct) k-mers. A set* 𝒴 = {*y*_1_, *y*_2_, … } *of m-mers is said to be a* multiminimizer cover *of* 𝒳 *if and only if, for any x* ∈ 𝒳, *there exists y* ∈ 𝒴 *and* 1 ≤ *i* ≤ *N, so that* min_*i*_(*x*) = *y, where* min_*i*_(*x*) *designates the smallest m-mer of x according to the order* 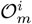. *In other words, for each k-mer x* ∈ 𝒳, *at least one of its N minimizer candidates belongs to* 𝒴.

The natural extension of the deduplicated density in this context would be 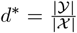, with the same trivial bounds as before. Contrary to MinCover_*f*_ (𝒳), we have here a choice of which candidate to choose as a minimizer, and therefore a chance at minimizing *d*^∗^ by making the appropriate choices. Therefore, this leads to considering the following problem.

*Problem 1 (M**ulti**M**in**C**over*). Provided a set 𝒳 of *k*-mers and *N* orders on *m*-mers 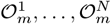, find a set 𝒴 of *m*-mers that is a multiminimizer cover of 𝒳 and of minimum cardinality |𝒴|.

This problem is closely related to the one of MinimumHIttingSet: provided an integer *k*, and a set 𝒳 of *k*-mers, find a set 𝒴 of *m*-mers (called an hitting set) such that each *k*-mer of 𝒳 contains at least one *m*-mer from 𝒴, and of minimum cardinality. This problem is known to be NP-hard [38,39]. Here the problem is slightly different, as we do not ask the elements of 𝒴 to simply appear in the *k*-mers of 𝒳, but also to be their minimizers for at least one of the orders. This problem also proves to be difficult, as per the following result.

#### Theorem 2.

*M**ulti**M**in**C**over**is NP-complete (for σ* ≥ 3 *and N* ≥ 2*). Proof*. The proof is deferred to Appendix S6.

Since minimizing |𝒴| is difficult, we must resort to heuristics. Furthermore, when parsing long sequences, it would be prohibitively expensive to keep track of all the *k*-mers encountered and their candidates minimizers before starting to build the multiminimizer cover. For this reason, we have developed a heuristic that makes decisions throughout the parsing of a sequence, based solely on the local context, that is presented in Section 3.3.

## 3 Results

In this section, we evaluate the performance of two implementations of the multiminimizer technique: an iterator over random-minimizers-based multiminimizers, and a proof of concept of multiminimizers using open-closed mod-minimizers [26]. Both are written in the Rust programming language, and freely available at github.com/lrobidou/multiminimizers. We propose results for canonical *k*-mers, but our implementation also supports non-canonical *k*-mers.

Density results are provided in Section 3.1, whereas the space usage is investigated in Section 3.2 and the conservation of sampled multiminimizers is studied in Appendix S8. Regarding running time, note that the multiminimizer scheme must compute all candidate minimizers, so iteration over multiminimizers is linear in the number of hash functions. We measured the time required to iterate over a random sequence and confirmed that our implementation scales linearly with the number of hash functions, while remaining very fast as shown in Appendix S9. Finally, we provide some results in Section 3.3 on multiminimizers applied to filtering approaches.

### 3.1 Density

As discussed in Section 2.3, the multiminimizer trick aims at reducing the density of any local scheme. We expect to reduce the density when increasing the number of hash functions. The intuition is that increasing the number of super-*k*-mer candidates increases the chance of finding a super-*k*-mer that ends further than the others, thus increasing the distance between consecutive selected positions.

To show this density reduction, we generated a random sequence (5M bases) and computed the density of our scheme on it, for 1 to 8 hash functions, using *w* = 15 and varying *m* (thus *k*). We observed that the higher the number of hash functions, the lower the density, which allows us to reach densities lower than any other scheme (see Figure 4a). To our knowledge, multiminimizer is the first iterator over minimizers with a density lower than the lower bound introduced by [27]. Figures for additional *m* and *w* values are available in the Appendix S7.

**Fig. 4:**
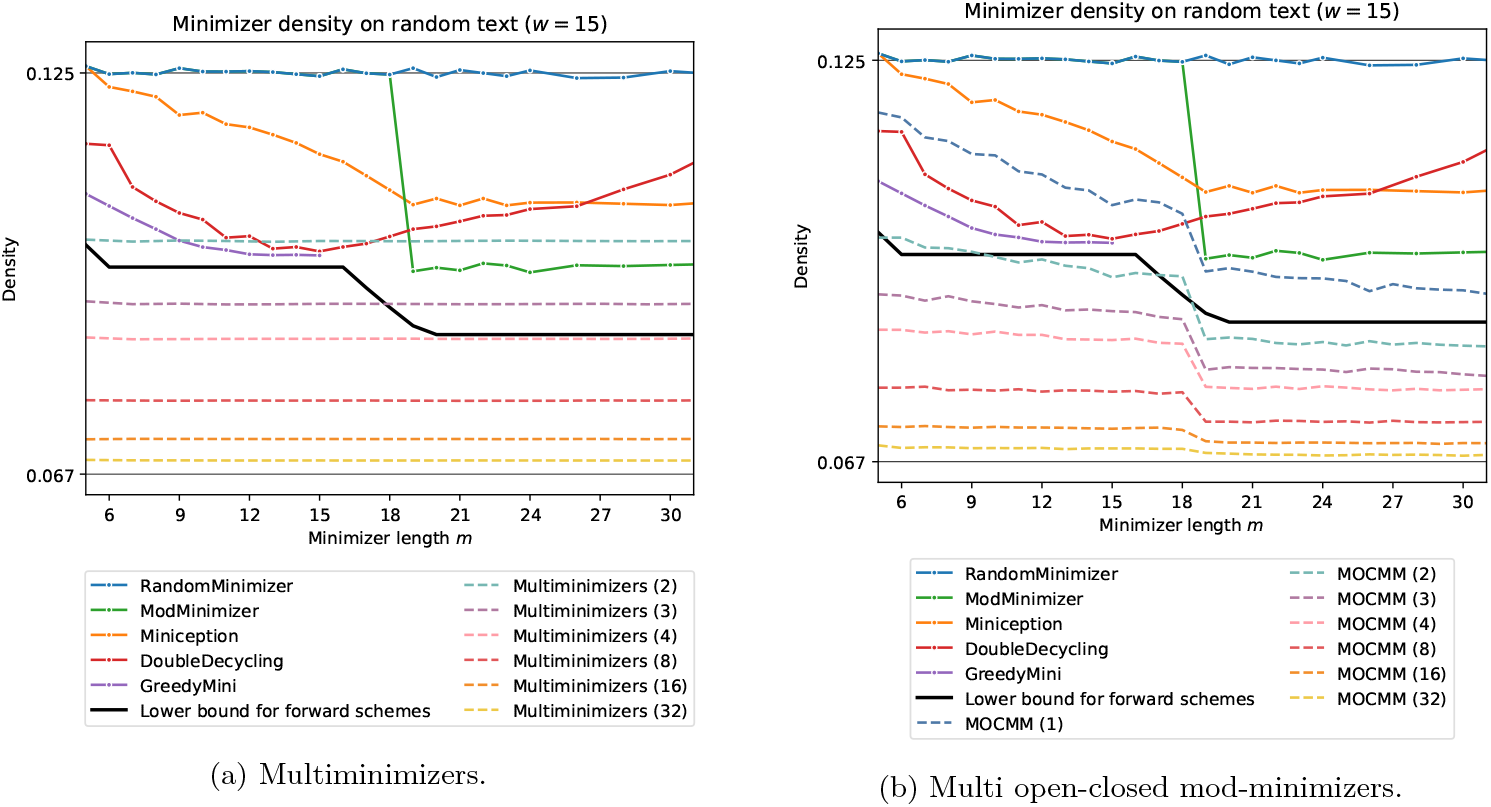
Density of multiple schemes, compared to our (a) multiminimizer and (b) MOCMM schemes with up to 32 hash functions. Our schemes are able to reach densities less than the lower bound of [27]. `GreedyMini` is close to this bound, but the computation time is exponential in *m* (the next data point would take ≈ 62h to compute).

Although multiminimizers are not a local scheme, we have observed that the empirical density 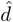 and the empirical average distance between selected positions 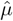 were indeed related as 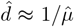, hence showing that Theorem 1 might apply beyond the scope of local schemes.

To show that our multiminimizer trick can be applied to other schemes than random minimizers, we applied it to the open-closed mod-minimizer [26] to obtain a proof-of-concept of a multi open-closed mod-minimizer scheme (MOCMM for short). Figure 4b shows that this novel scheme achieves a lower density than the multiminimizers on a random sequence. At the time of writing, the library we used to compute the minimizer scheme does not support the open-closed mod-minimizer scheme. Therefore, we used a non-SIMD implementation, making it much slower. For this reason, we could only reasonably run it on a random sequence of 5 *×* 10^5^ bases.

In both Figure 4a and Figure 4b, one can observe that the density approaches the theoretical lower bound 1*/w*. This convergence seems intuitive since, when the number of hash functions becomes large enough, all *m*-mers of a *k*-mer are likely to be chosen as the minimizer candidate for at least one of the hash functions, and therefore the multiminimizer scheme tends to become optimal.

### 3.2 Super-*k*-mers and hyper-*k*-mers space usage

Since the space usage of a super-*k*-mer representation of a sequence is driven by the density of the underlying scheme [17, Thm. 1], we can translate a density reduction to a linear super-*k*-mer representation. Figure 5a shows the space usage of a super-*k*-mer representation of multiminizers, whereas Figure 5b provides the same information for MOCMMs, both for multiple numbers of hash functions and *k* values.

**Fig. 5:**
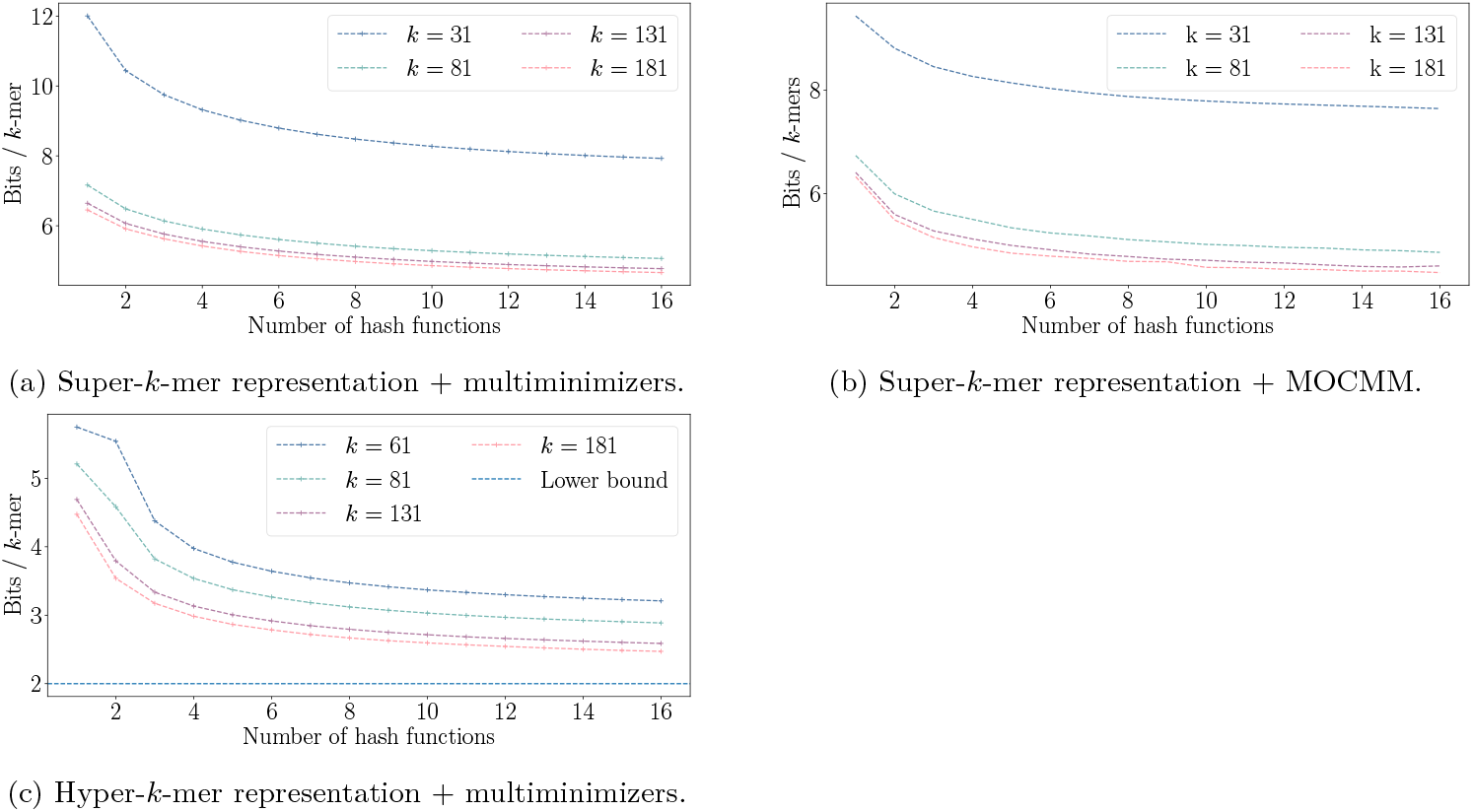
Number of bits per *k*-mer for different representations of a random read, with *m* = 21, depending on *k* and the number of hash functions used with the multiminizer trick. Increasing the number of hash functions allows the hyper-*k*-mer representation (c) to converge to 2 bits/*k*-mer, the minimum required to represent a DNA sequence.

Ideally, to represent a sequence over an alphabet of size 4 (e.g., a DNA sequence *S*), one could use only 2 bits per nucleotide in *S*. However, super-*k*-mers use as low as 6 bits per nucleotide in *S*. Hyper-*k*-mers [17] were recently introduced and tend toward 4 bits per nucleotide in *S* when using a random-minimizer-based hyper-*k*-mer representation. This is still higher than the lower bound of 2 bits per nucleotide, due to random minimizers having a density of 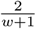.

Since our multiminimizer scheme approaches a density of 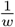, using a multiminimizer scheme could allow hyper-*k*-mers to tend toward 2 bits per nucleotide in the input sequence. To test this hypothesis, we replaced the iterator over minimizer of KFC (a hyper-*k*-mer-based *k*-mer counter introduced in [17]) by our iterator over multiminimizer and report the result in Figure 5c. Note that KFC is optimized for large *k* values (e.g. *>* 60), so we tested our hypothesis in this range. Our results show that KFC converges to 2 bits per base in *S*. To the best of our knowledge, this is the first streaming *k*-mer representation to achieve it.

### 3.3 Pin: a multiminimizer approach to Needle-like applications

We developed a simple proof-of-concept minimizer index to demonstrate the utility of the multiminimizers approach for Needle-like filtering stages, in a prototype dubbed Pin (github.com/Malfoy/Pin). Such an index stores a set of minimizers that collectively cover all *k*-mers of interest, enabling efficient filtering as an indexed minimizer is a necessary condition for an indexed *k*-mer, and with sufficiently large minimizers, passing the filter already implies substantial sequence similarity. Our prototype relies on exact hash tables and handles a single uncolored set; the same idea can be readily adapted to colored or approximate index structures. During construction, we test the presence of *k*-mers and then greedily select minimizers to cover any uncovered sequences. We evaluate the approach on the Human HiFi accession `SRR11292123` (24Gb) in Figure 6, and observe a substantial reduction in index size with competitive build and query times. Merely switching from a single to two hash functions yields an index about 20% smaller, with roughly a 20% increase in construction time and about 85% in query time.

**Fig. 6:**
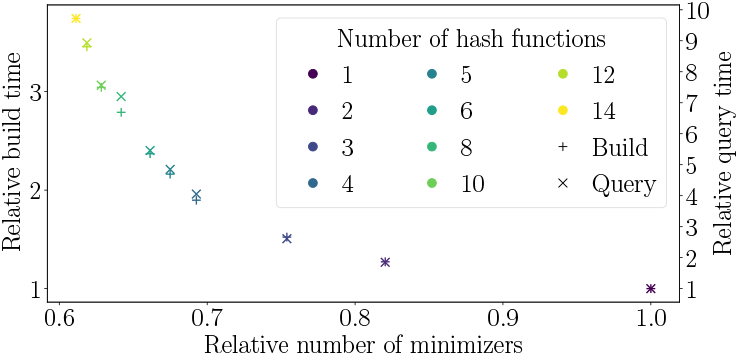
Relative performances of the minimizer dictionary Pin using multiminimizers, comparing *N* hash functions vs *N* = 1. *x*-axis depicts the relative index size (in terms of the number of minimizers), and *y*-axis provides relative construction (left) and query (right) times. As an indicator, for *N* = 1, there are ≈ 850 *×* 10^6^ minimizers, construction time is 234s, and query time is about 0.04s.

## 4 Future work

This work advances both the theory and practice of minimizer-based sampling by (i) establishing a general density–distance equivalence under minimal assumptions (**??**); (ii) introducing the novel notion of the deduplicated density, which captures the fraction of distinct minimizers covering a set of *k*-mers and for which we provide an analytical expression and empirical characterization in the random minimizers case; and (iii) designing the multiminimizer scheme, a simple yet effective scheme that trades time for space and achieves densities below the forward-scheme lower bound while preserving linear runtime. We also propose a proof-of-concept of a multi open-closed mod-minimizer scheme, achieving even lower densities. Empirically, increasing the number of hash functions consistently reduces density and memory footprint, and the benefits translate directly to super-*k*-mers, hyper-*k*-mers representations, and filtering indexes.

Several theoretical questions remain open. First of all, the theoretical density of the multiminimizer scheme remains to be studied. Since it empirically reaches densities lower than the bound for local schemes [27], it would be of great interest to find a way to express its density both in terms of the number and the underlying densities of the schemes used to build the multiminimizer meta scheme. Additionally, theoretical guarantees on the conservation of multiminimizers remain to be established. Then, the computational complexity of globally minimizing the standard density is still unknown, despite our NP-completeness result for the deduplicated counterpart; clarifying this complexity, including connections to polar-set formulations, is a natural next step. Beyond random minimizers, extending the analysis of deduplicated density to other scheme families and simplifying its expression in the random case would strengthen the theoretical picture. From an algorithmic standpoint, developing effective local or global heuristics for MultiMinCoveris a promising path. Finally, on the practical side, two areas can be explored. Firstly, combining multiminimizers with other existing low-density schemes to form multi-seeded variants could further reduce density in practice while retaining fast iteration. The proof of concept “multiminimizers on open-closed mod-minimizer” could be vastly improved by, e.g., using SIMD instructions to speed up computation. Secondly, replacing traditional minimizer iterators with our Rust library could improve a vast number of existing tools, beyond our work here on improving a hyper-*k*-mer-based *k*-mer counter. Thus, we set the stage for the next generation of theoretically grounded, robust, and efficient sampling schemes. Different sequence-mapping tools have highlighted the need for heuristics to fill in the deserts where sampled alignment landmarks are missing, as well as to better handle low complexity regions. We believe that the formalization of choice and selection-based schemes, such as those introduced in this work, will support improved alignment strategies in future developments.

## Acknowledgments

This work was supported by the French National Research Agency AGATE [ANR-21-CE45-0012] and full-RNA [ANR-22-CE45-0007]. With financial support from ITMO Cancer of Aviesan within the framework of the 2021-2030 Cancer Control Strategy, on funds administered by Inserm. The authors are grateful to three anonymous reviewers for their valuable comments on a first version of the manuscript.

## Supplementary material

### S1 Proof of Theorem 1

We start by rewriting (1) as

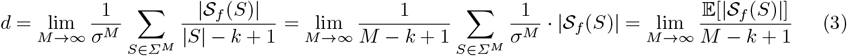

where 𝔼[|𝒮_*f*_ (*S*)|] designates the expected particular density of a (uniform) random sequence *S* of length *M*, seen as a random variable. Suppose we can extend the original sequence *S*, of size *M*, and continue selecting minimizers past the first *M* characters. We define 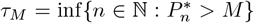 as the random variable that determines how many minimizers have been selected in *S*; hence *τ*_*M*_ = |𝒮_*f*_ (*S*)| . Therefore, we can rewrite (3) as 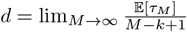.

Let us call 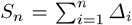. Notice that, by definition of the 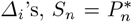; then *τ*_*M*_ = inf*{n* ∈ ℕ : *S*_*n*_ *> M*} is a stopping time with respect to the *σ*-algebra generated by the Δ_*i*_’s. Using Wald’s equation [40], we get

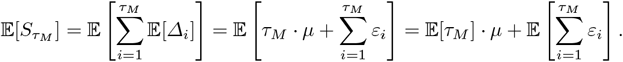

We apply again Wald’s equation to the right-hand term and obtain

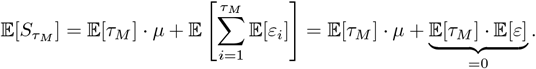

By definition of *S*_*n*_, *τ*_*M*_ and the Δ_*i*_’s, notice that 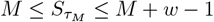. Therefore

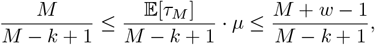

and, taking the limit as *M* → ∞, we establish our result.

### S2 Proof of Proposition 1

We begin with the following lemma.

#### Lemma 2.

*For any function f* : [0, 1] → ℝ *at least two times continuously derivable, we have*

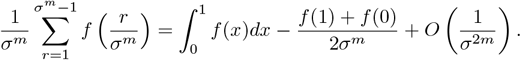

*Proof*. This is a direct application of the trapezoid rule for numerical integration. Defining 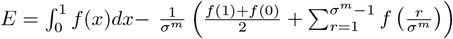 we have 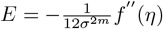 for some *η* ∈ [0, 1] [41, eq. (5.1.17)].

Let 1 ≤ *i* ≤ *w*. For *R*_*i*_ to be the minimizer of the window *R*_1_, …, *R*_*w*_, then we must have the following situation :

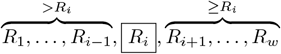

The events (*R*_*j*_ *> R*_*i*_) and (*R*_*j*_ ≥ *R*_*i*_) are clearly not independent of *R*_*i*_; but resorting to the law of total probability, and that the *R*_*j*_’s are i.i.d., we obtain

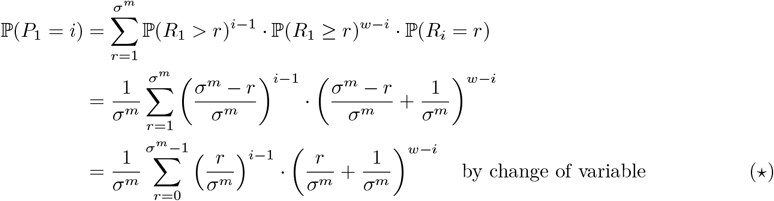

Using Lemma 2, with *f* : *x 1*→ *x*^*i*−1^(*x* + *σ*^−*m*^)^*w*−*i*^, we obtain

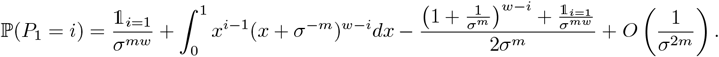

Using the fact that 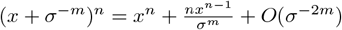, we get

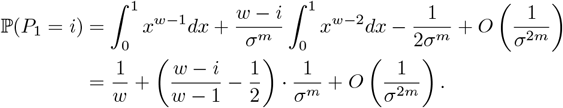

### S3 Convergence of *P*_1_ to the uniform distribution

In Figure 7, we computed the distribution of *P*_1_ for various values of *m*, using the exact value stated in ( ) rather than the approximation of Proposition 1. As one can see, *P*_1_ converges quickly to the limit uniform distribution (as one can expect since the dependency on *m* is of the order of *σ*^−*m*^).

**Fig. 7:**
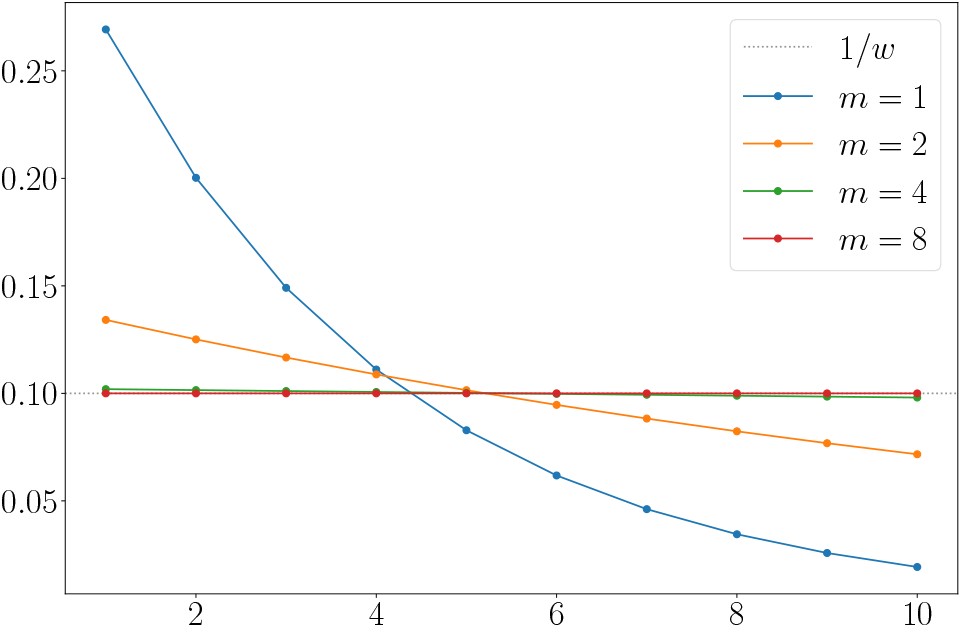
With *w* = 10, evolution of the distribution of *P*_1_ as *m* increases. The actual value of *P*_1_ is computed, using ( ) instead of the approximations of Proposition 1.

### S4 Monte-Carlo simulations of random minimizers

Simulating our probabilistic model of random minimizers, with Hypothesis 2, we obtain the results depicted in Figure 8, which have been commented on in the core of the paper.

**Fig. 8:**
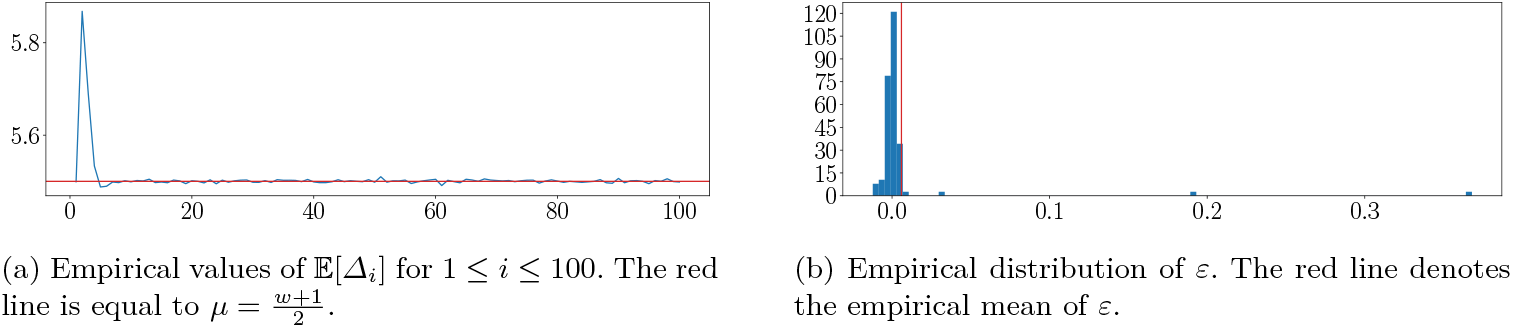
Monte-Carlo simulations of our model of random minimizers under Hypothesis 2. We used *m* = 8, *w* = 10 and generated *N*_simu_ = 10^6^ random sequences **R**, all long enough so that the sequence **Δ** = Δ_1_, Δ_2_, … contains at least 100 values.

One thing that seems surprising at first glance is the peak observed for 𝔼[Δ_2_]. When sliding the window by one position, either the previous minimizer *X* is still in the window, and we change the minimizer only if the new value *X*^′^ is *< X*, or either *X* is not in the window anymore, and we select as a new minimizer the minimum value in the window (which is called a *rescan*). Here, *P*_1_ is always obtained by a rescan operation, meaning that 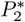 is always obtained as “the next minimizer following a rescan”. We do not observe any other peak in the distribution of 𝔼[Δ_*i*_]’s as the other rescans are randomly found in the sequences, and therefore their impact is smoothed among all simulations. We have the following result; note that both the formula and the empirical simulations yield a value of 𝔼[Δ_2_] ≈ 5.87 for *w* = 10 and *m* = 8.

#### Proposition 3.

*Assuming* 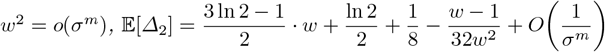.

We actually prove the following, more general, result:

#### Proposition 4.

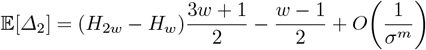

*where H*_*n*_ *denotes the n-th harmonic number* 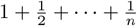.

*Proof*. To prove this, we need to distinguish two cases: either 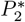 “overwrites” 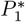 because 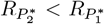 or 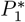 goes out of scope and 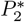 is selected by a rescan. Therefore, we have

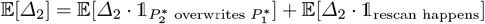

In the following paragraphs, we show that

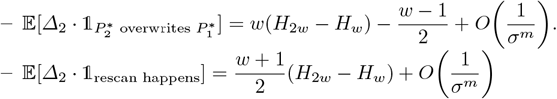

Finally,

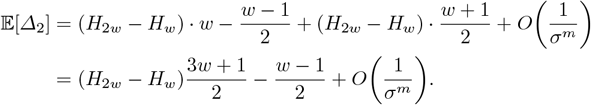

By approximating *H*_*n*_ as ln 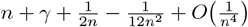, we get the desired result:

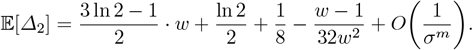

*Overwrite case*. We start with the following lemma.

#### Lemma 3.

*For any* 1 ≤ *j* ≤ *i* ≤ *w, the probability that* 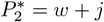 *overwrites* 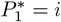 *is*

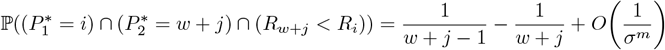

*Proof*. Let *O*_*i,w*+*j*_ denote the event 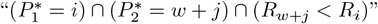,

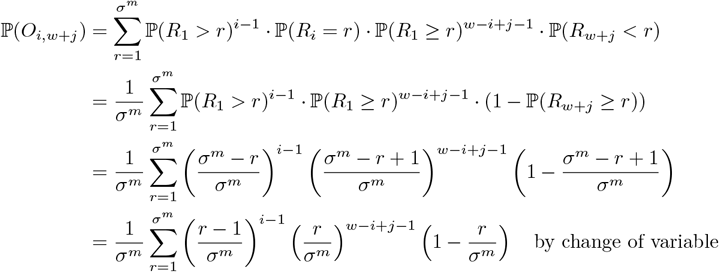

Using Lemma 2 with *f* : *x 1*→ (*x* − *σ*^−*m*^)^*i*−1^*x*^*w*−*i*+*j*−1^(1 − *x*), we obtain

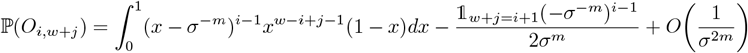

Using the expansion (*x* − *σ*^−*m*^)^*i*−1^ = *x*^*i*−1^ + *O*(*σ*^−*m*^), we get

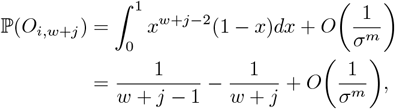

since 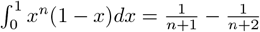.

Assuming *w*^2^ = *o*(*σ*^*m*^),

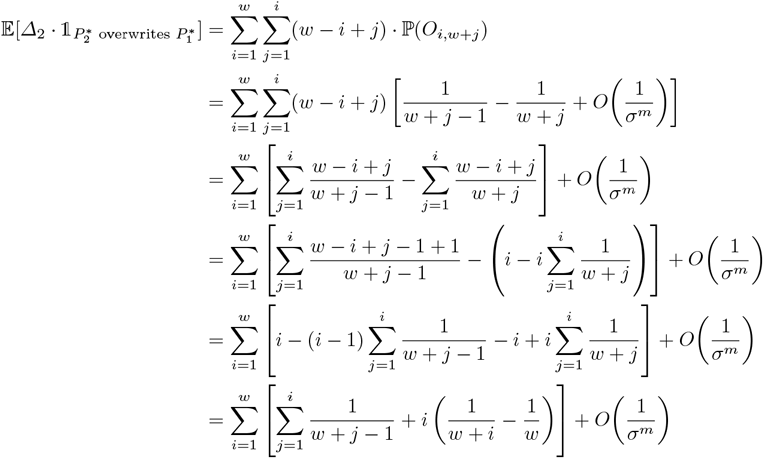

Expanding into three different sums, we have

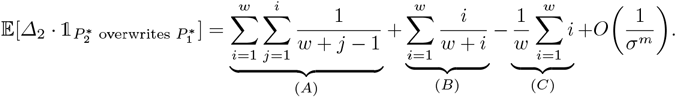

(*C*) is obviously 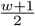. For (*A*), note that the term 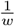 appear *w* times, 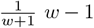 times, and so on up to 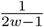 that appear only once. Hence,

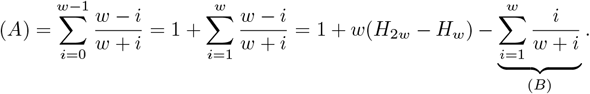

We do not need to compute (*B*) as terms cancel each other out, and

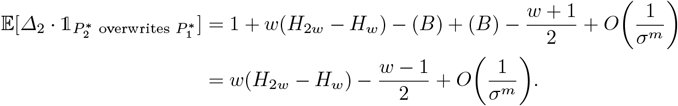

*Rescan case*. Let 1 ≤ *r* ≤ *σ*^*m*^ and let 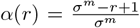. We use the trapezoid rule used in Lemma 2 to derive an alternative formula:

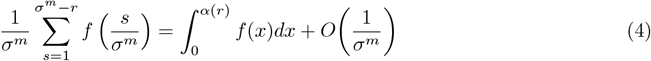

where *f* : [0, 1] → ℝ is twice continuously derivable.

Then, we have the following lemma.

#### Lemma 4.

*For any* 1 ≤ *i, j* ≤ *w, the probability that* 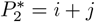 *as a rescan when* 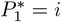 *is*

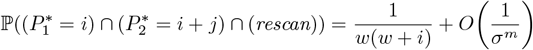

*Proof*. Let us denote by *R*_*i,j*_ the event 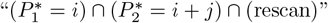 for any 1 ≤ *i, j* ≤ *w*. When this event is realized, then all variables

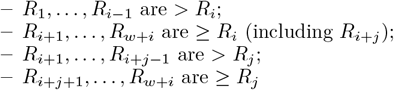

Therefore,

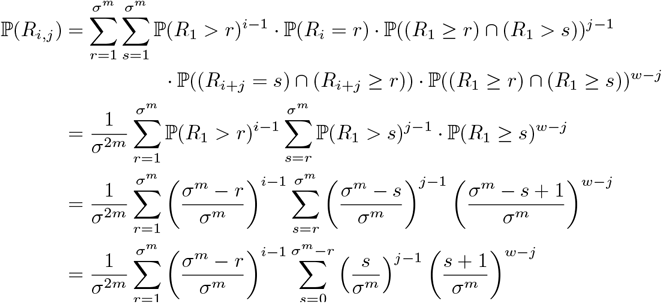

Notice that the term *s* = 0 in the inner sum equals 𝟙_*j*=1_*σ*^−*m(w*−1)^ which is *O*(*σ*^−*m*^) as we suppose that *w >* 1. Applying (4) to the function *f* : *x 1*→ *x*^*j*−1^(*x* + *σ*^−*m*^)^*w*−*j*^, we get

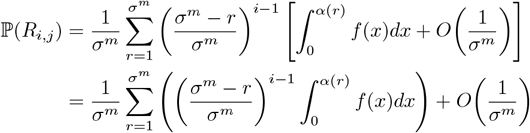

Using the expansion *f* (*x*) = *x*^*w*−1^ + *O*(*σ*^−*m*^), we get

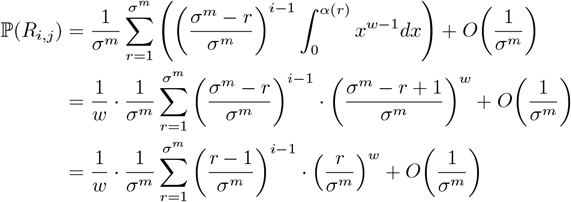

Applying Lemma 2 with the function *f* : *x 1*→ (*x* − *σ*^−*m*^)^*i*−1^*x*^*w*^, we get

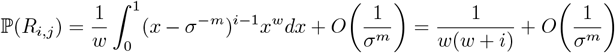

Finally, still assuming that *w*^2^ = *o*(*σ*^*m*^)

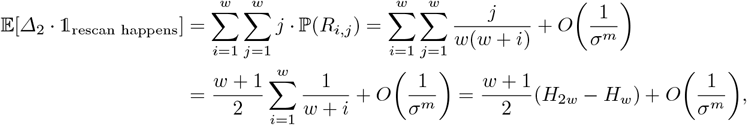

as claimed.

### S5 Proof of Proposition 2

We are interested here in computing the deduplicated density *d*^∗^ of the random minimizer scheme. We have, from (2),

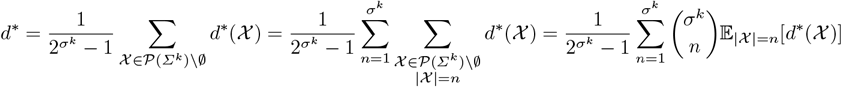

To compute the required expectation, let 𝒳 = {*X*_1_, …, *X*_*n*_*}* be a set of *k*-mers, chosen uniformly at random. We denote by *M*_*i*_ ∈ [[1, *σ*^*m*^ ] the random variable denoting the minimizer of *k*-mer *X*_*i*_ (assimilating minimizers and their rank according to the chosen order 𝒪_*m*_). Under Hypothesis 2, it is not difficult to establish that ℙ (*M*_*i*_ *> r*) = ℙ(*X*_1_ *> r*)^*w*^ where *X*_*i*_ denotes the *i*-th *m*-mer of *X*_*i*_ and *w* = *k* − *m* + 1. Therefore, we obtain

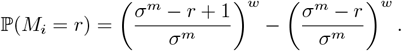

We denote by *P*_*r*_ the number of *k*-mers in 𝒳 that admits *r* as their minimizers; and we are interested in computing

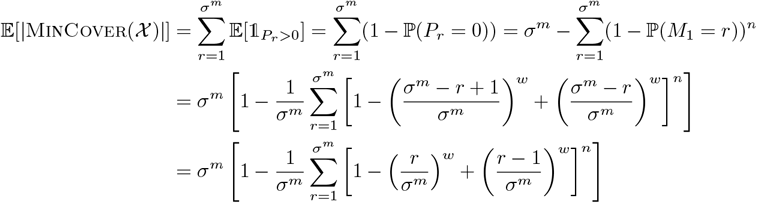

This is exactly the standard coupon collector’s problem, where the coupons are distinct minimizers and |𝒳| is the number of coupons bought.

### S6 Proof of Theorem 2

We prove the NP-completeness of MultiMinCover by (i) showing first that the problem is in NP and then (ii) by reduction from SetCover. One anonymous reviewer also suggested an alternative reduction from VertexCover, which is presented at the end of this section.

#### MultiMinCover is in NP

For this, we look at the decision version of MultiMinCover. Provided an integer *k*, a set 𝒳 of *k*-mers and *N* orders on *m*-mers 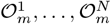, is there a set 𝒴 of *m*-mers that is a multiminimizer cover of 𝒳 and with cardinality |𝒴| ≤ *k*? If one provides a set ᵋ of *m*-mers of cardinality ≤ *k*, one can trivially check whether 𝒴 is a multiminimizer cover of 𝒳 by simply iterating over *k*-mers in 𝒳, compute their *N* minimizers and check whether at least one of them belong to 𝒴. This can be done in *O*(|𝒳| · *N* ), hence the result.

*Reduction from SetCover*. Let 𝒰 = *{*1, …, *n}* and let 𝒮 ⊆ 2^*U*^ be a collection of subsets of 𝒰, whose union is 𝒰 . We recall that the SetCover problem amounts to finding the smallest subcollection of 𝒮 whose union is 𝒰 . The *frequency* of an element *u* ∈ 𝒰 is defined at the number of subsets in 𝒮 that contains *u*. We define *N* = max_*u*∈𝒰_ freq(*u*) as the maximum frequency of any element of 𝒰 . Note that *N* ≤ | 𝒮 |.

*Example 1*. To illustrate the proof, we propose to follow an example. Let 𝒰 = {1, …, 5} and 𝒮 = {{1, 2, 3}, {4, 5}, {2, 4},{3, 4}}. The optimal solution for this instance of set cover is {1, 2, 3}, {4, 5} of cardinality 2. The maximal frequency is *N* = 3 (since 4 appears in 3 subsets).

We use the following 3-letter alphabet Σ = {0, 1, *a}* . Let *m* = ⌈ log_2_ |𝒮|⌉. Take any indexing *S*_1_, …, *S*_|𝒮|_ of the subsets forming 𝒮; we associate to each subset *S*_*i*_ the *m*-mer *P*_*i*_ corresponding to *i* in binary coded with *m* bits. Then, let *k* = *Nm*+*N* − 1 = *N* (*m*+1) − 1. To each element *u* ∈ 𝒰, let *i*_1_ …, *i*_*f*_ be the indices of the *S*_*i*_’s containing *u*, with *f* = freq(*u*). We associate to *u* the *k*-mer 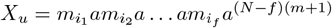.

*Example 2*. We have *m* = ⌈log_2_ 4⌈ = 2. We associate *m*_1_ = 00 to {1, 2, 3*}, m*_2_ = 01 to {4, 5*}, m*_3_ = 10 to {2, 4*}* and *m*_4_ = 11 to {3, 4*}*. We have *k* = 8 and the following *k*-mers

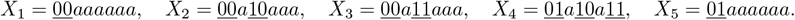

The goal now is to exhibit *N* orders 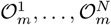 on *m*-mers, so that the following is true: for each *k*-mer *x*, its *N* minimizers correspond exactly to its *m*-mers coming from the *m*_*i*_’s; i.e. its minimizers contain only the letters 0 and 1. It should be clear that any solution to MultiMinCover in this context is equivalent to a solution of the original SetCover instance. Since there are |𝒰| = *n k*-mers of size *N* (*m* + 1) − 1 = *O*(|𝒮| log_2_ |𝒮|) and exactly |𝒮| minimizers to chose from, the reduction is indeed polynomial, hence the result — once we prove said orders on *m*-mers can indeed be constructed in polynomial time.

By construction, we want the *m*-mers *m*_1_, …, *m*_|𝒮|_ to receive a rank (for all orders) between 1 and |𝒮|; all subsequent possible *m*-mers (either binary, or containing at least one ‘*a*’) should receive a rank *>* |𝒮| + 1, and their relative order is of no importance since they will not be selected as minimizers whatsoever. By doing so, we ensure that the *N* minimizers of each *k*-mer *X*_*u*_ can only be the *m*_*i*_’s it is made of. If *X*_*u*_ contains several such *m*_*i*_’s, the only thing left to do is to ensure that each of them is indeed a minimizer for at least one order – so that the number of distinct minimizers of *X*_*u*_ is equal to the frequency of *u* in the original SetCover instance — thus ensuring the 1-to-1 correspondence between solutions of the two problems.

We exhibit Algorithm 2 to construct those orders.

*Example 3*. At this point, we have determined that any *m*-mer containing at least one ‘*a*’ has rank at least 5 in any of the 3 orderings to build, and {00, 01, 10, 11} must receive ranks 1 to 4. We initialize two arrays, `ranks` and `minimizers`, following Algorithm 2. Applying Lines 7-9 to *X*_1_ and *X*_5_, we get:

| rank | $\mathcal{O}_m^1$ | $\mathcal{O}_m^2$ | $\mathcal{O}_m^3$ | minimizers | $\min_1(x)$ | $\min_2(x)$ | $\min_3(x)$ | cand(x) |
| --- | --- | --- | --- | --- | --- | --- | --- | --- |
| 00 | | | | $X_1$ | 00 | 00 | 00 | ∅ |
| 01 | | | | $X_2$ | | | | 00, 10 |
| 10 | | | | $X_3$ | | | | 00, 11 |
| 11 | | | | $X_4$ | | | | 01, 10, 11 |
| | | | | $X_5$ | 01 | 01 | 01 | ∅ |

Now, consider *X*_2_ and the *m*-mer 00. Applying Lines 11-15, choosing *l* = 1, we get:

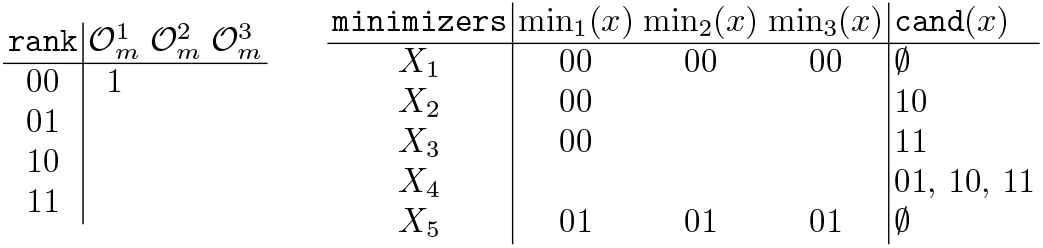

Now, consider $X_2$ and the $m$ -mer 00. Applying Lines 11-15, choosing $l = 1$ , we get:
| rank | $\mathcal{O}_m^1$ | $\mathcal{O}_m^2$ | $\mathcal{O}_m^3$ | minimizers | $\min_1(x)$ | $\min_2(x)$ | $\min_3(x)$ | cand(x) |
| --- | --- | --- | --- | --- | --- | --- | --- | --- |
| 00 | 1 | | | $X_1$ | 00 | 00 | 00 | ∅ |
| 01 | | | | $X_2$ | 00 | | | 10 |
| 10 | | | | $X_3$ | 00 | | | 11 |
| 11 | | | | $X_4$ | | | | 01, 10, 11 |
| | | | | $X_5$ | 01 | 01 | 01 | ∅ |

For *X*_2_ and *X*_3_, their respective remaining candidate fill the empty spots in `minimizers` by applying again Lines 7-9. Considering *X*_4_, a possible outcome could be the following:

| rank | $\mathcal{O}_m^1$ | $\mathcal{O}_m^2$ | $\mathcal{O}_m^3$ | minimizers | $\min_1(x)$ | $\min_2(x)$ | $\min_3(x)$ | cand(x) |
| --- | --- | --- | --- | --- | --- | --- | --- | --- |
| 00 | 1 | | | $X_1$ | 00 | 00 | 00 | ∅ |
| 01 | 2 | | | $X_2$ | 00 | 10 | 10 | ∅ |
| 10 | | 1 | | $X_3$ | 00 | 11 | 11 | ∅ |
| 11 | | | | $X_4$ | 01 | 10 | 11 | ∅ |
| | | | | $X_5$ | 01 | 01 | 01 | ∅ |

After which, any completion of `ranks` would work.

##### Algorithm 2

DetermineOrders

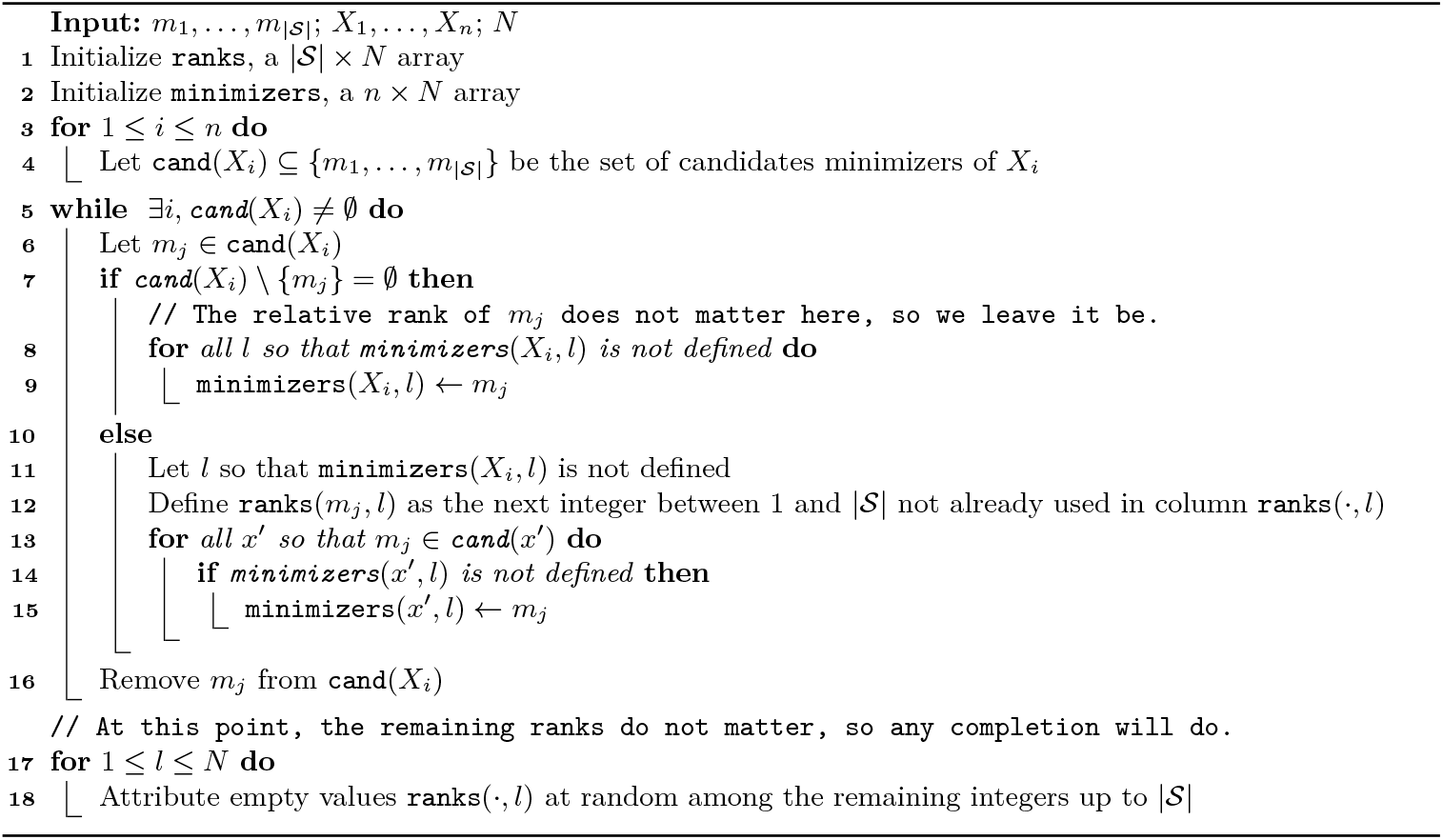

##### Proposition 5.

*Algorithm 2 is correct and runs in polynomial time*.

*Proof*. Each iteration in the `while` loop eliminates one candidate minimizer, of which there are at most |𝒰|· *N* . For a given candidate minimizer, if Lines 7-9 are executed, the `for` loop takes at most *N* steps; and if Lines 11-15 are executed, parsing a column in takes |𝒮| steps, and the loop at most |𝒰| steps. Since *N* ≤ |𝒮|, we end up with a worst-case complexity of *O* |𝒰| · |𝒮| · (|𝒮| + |𝒰|), polynomial as claimed.

For the algorithm to be correct, we must never encounter the case where there exists *X*_*i*_ and *m*_*j*_ ∈ `cand`(*X*_*i*_) such that `minimizers`(*X*_*i*_, *l*) is already defined for all 1 ≤ *l* ≤ *N* . In such a case, *m*_*j*_ could not be a minimizer for *X*_*i*_. Suppose we are in such a case. Let `cand`_*I*_ (*X*_*i*_) be the set of candidates of *X*_*i*_ at the initialization of the algorithm. Since *m*_*j*_ ∈ `cand`_*I*_ (*X*_*i*_) and since `minimizers`(*X*_*i*_, *l*) is defined for all *l*, then surely there exists *m*_*j*_′ ∈ `cand`_*I*_ (*X*_*i*_), *j*^′^ ≠ *j*, that appears at least twice in the list of minimizers of *X*_*i*_, by pigeonhole principle ( |`cand`_*I*_ (*X*_*i*_) \{*m*_*j*_}| *< N* ). When a minimizer is affected as a consequence of Lines 11-15, it is removed from the list of candidates (and therefore cannot be added again as a minimizer for another order), so the only way *m*_*j*_′ could have been affected to two orders if as a consequence of Lines 7-9 — but there are only triggered if *m*_*j*_′ is the only remaining element of `cand`(*X*_*i*_), thus a contradiction since it still contain at least *m*_*j*_.

*Reduction from V**ertex**C**over*. This alternative reduction was proposed by an anonymous reviewer. Let *G* = (*V, E*) be graph. Using the same formalism as above, associate to each vertex *i* a distinct minimizer *m*_*i*_ = `binary`(*i*) encoded with *m* = ⌈|*V*|⌉ bits. As above, we use the alphabet Σ = {0, 1, *a*} .

For each edge (*i, j*) ∈ *E*, assume w.l.o.g *m*_*i*_ *< m*_*j*_, we construct the *k*-mer *X*_(*i,j*)_ = *m*_*i*_*am*_*j*_. We define two orders 𝒪^1^ and 𝒪^2^ as (*m*_*i*_, *m*_*j*_) ∈ 𝒪^1^ ⇔ *m*_*i*_ *< m*_*j*_ and (*m*_*i*_, *m*_*j*_) ∈ 𝒪^2^ ⇔ *m*_*i*_ *> m*_*j*_. Finally, we complete the orders by saying that *x < y* if *a* appears in *y* and not in *x*, for both orders 𝒪^1^ and 𝒪^2^. If *a* belong to both *x* and *y*, any arbitrary order can do. In the end, for all *k*-mers *X*_(*i,j*)_, min_1_(*X*_(*i,j*)_) = *m*_*i*_ and min_1_(*X*_(*i,j*)_) = *m*_*j*_. Let 𝒳 = *{X*_(*i,j*)_ : *{i, j}* ∈ *E, m*_*i*_ *< m*_*j*_*}*.

##### Lemma 5.

*V**ertex* *C**over* *admits a yes-instance* (*V, E, k*) *iff MultiMinCover instance* ( 𝒳, 𝒪^1^, 𝒪^2^, *k*) *is a yes-instance*.

*Proof*. (⇒) If (*V, E, k*) is a yes-instance with vertex cover 𝒞, then the set 𝒴 = {*m*_*i*_ : *I* ∈ 𝒞} is a multiminizer cover of size *k* for 𝒳 since for any *k*-mer *X*_(*i,j*)_ either min_1_(*X*_(*i,j*)_) = *m*_*i*_ or min_2_(*X*_(*i,j*)_) = *m*_*j*_.

(⇐) Let (*V, E, k*) be a no-instance. Assume that our constructed instance (𝒳, 𝒪^1^, 𝒪^2^, *k*) is a yes-instance with multiminimizer cover 𝒴. Then every *X*_(*i,j*)_ ∈ 𝒳 has a minimizer min_1_(*X*_(*i,j*)_) = *m*_*i*_ or min_2_(*X*_(*i,j*)_) = *m*_*j*_ per definition of 𝒪^1^ and 𝒪^2^; and the set 𝒞 = *{i* : *m*_*i*_ ∈ 𝒴*}* is a vertex cover of size *k* for *G* = (*V, E*) with *E* = {(*i, j*) : *X*_(*i,j*)_ ∈ 𝒳*}*.

### S7 Density plots for multiple *w* values

In this section, we ran the same evaluation as in Section 3.1, but applied our multiminimizer iterator to multiple *w* and more *m* values to check that our results are robust with regard to *w* and *m*. Figure 9 and Figure 10 shows that our results are indeed robust with regard to *w*: for all *w* values, our iterator can yield densities lower than the lower bound.

**Fig. 9:**
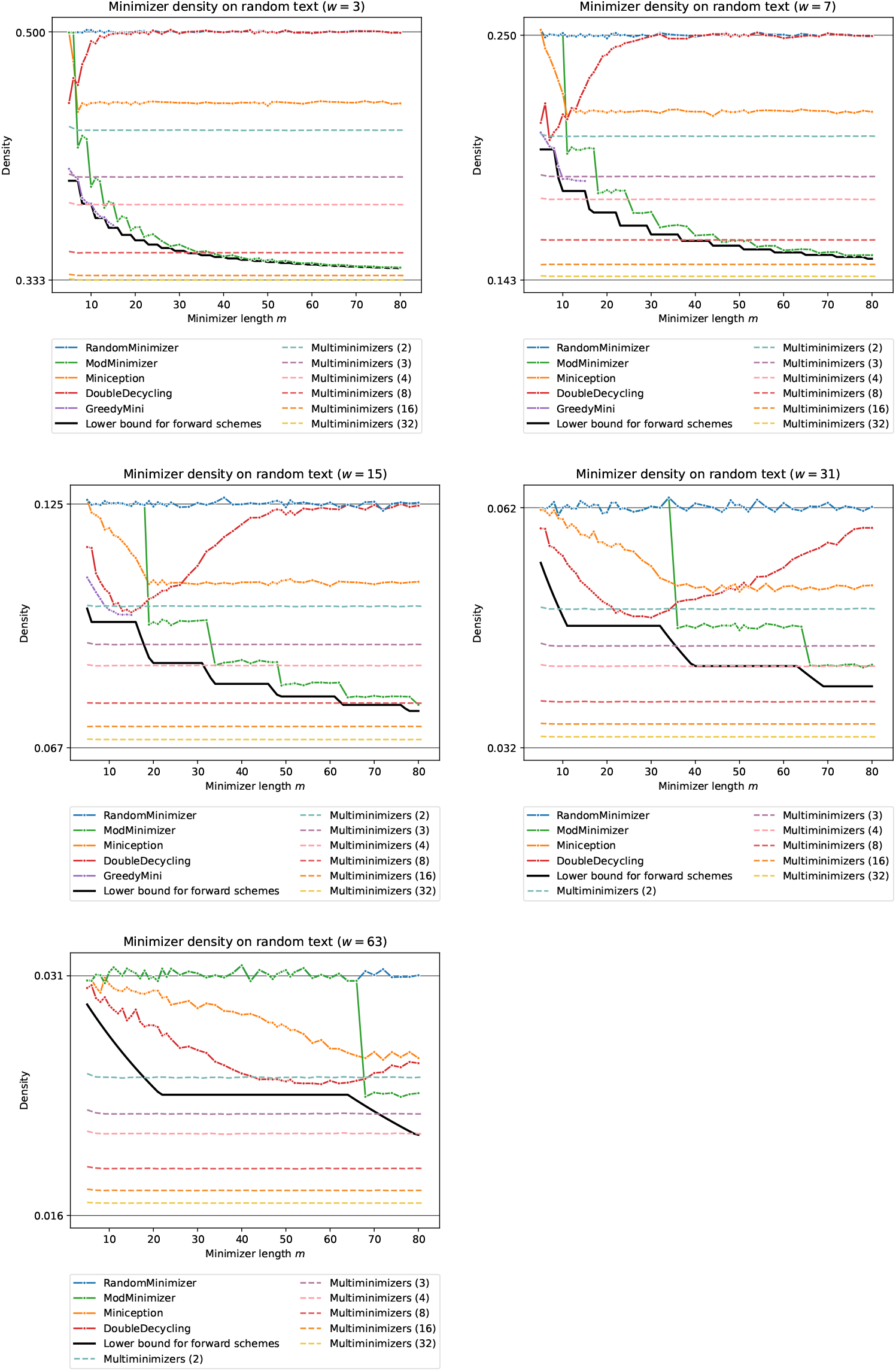
Density of multiple schemes, including multiminimizers with up to 32 hash functions. Same plot as Figure 4a, for *w* ∈ {3, 7, 31, 63*}*.

**Fig. 10:**
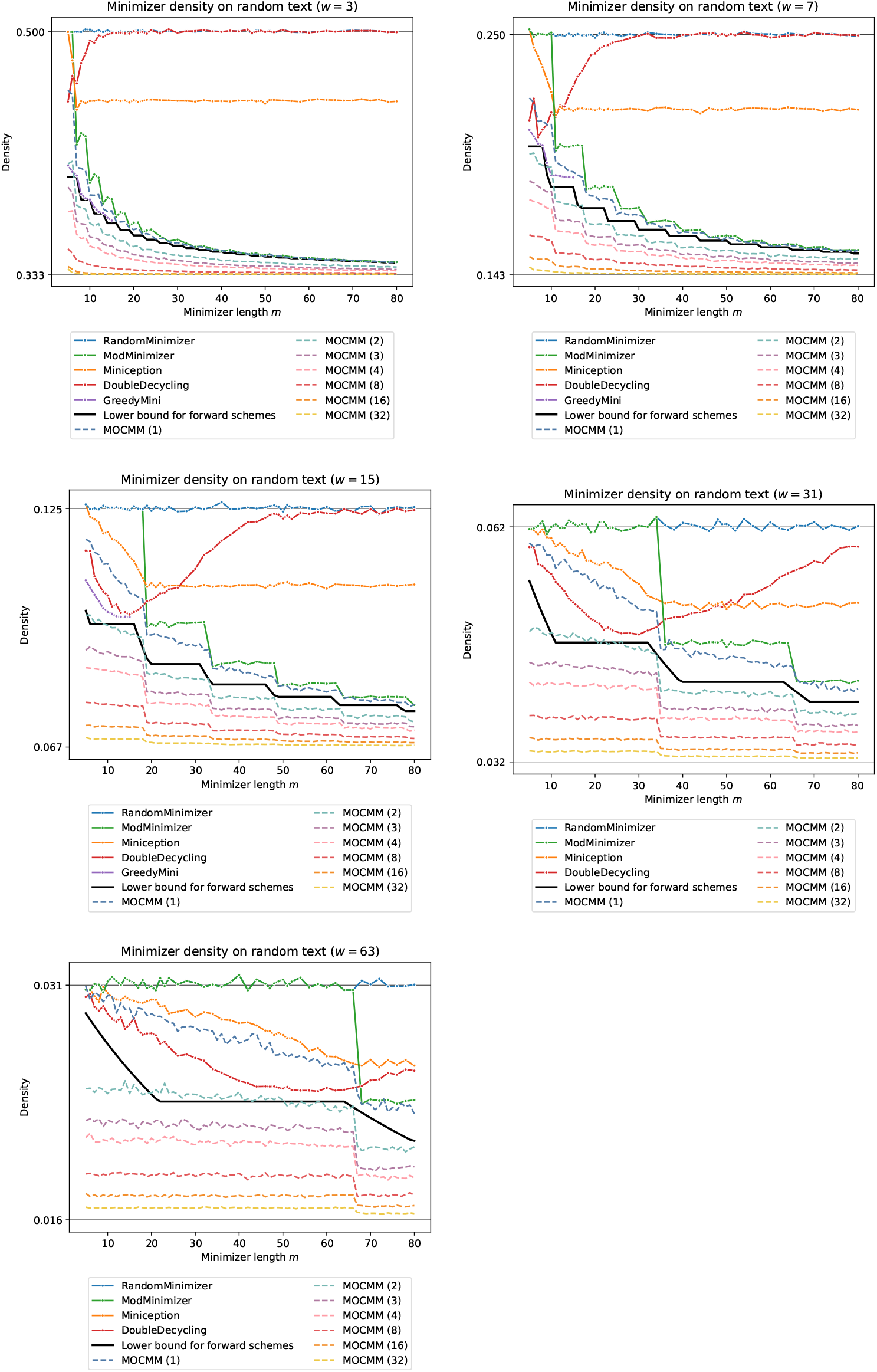
Density of multiple schemes, including multiminimizers based on open-closed mod-minimizers [26] with up to 32 hash functions. Same plot as Figure 4b, for *w* ∈ {3, 7, 31, 63*}*.

### S8 Conservation with respect to error rate

In this section, we study the evolution of the conservation of the sampled multiminimizers with respect to the error rate and the number of hash functions. Given a sequence *S* and a mutated version of the sequence *S*^′^ for a given error rate, we measure the conservation using the Jaccard similarity of *A* = `MultiMinimizers`(*S*) and *B* = `MultiMinimizers`(*S*^′^), i.e. 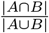. The results are shown in Figure 11. In particular, we can observe that the conservation decreases slightly faster with respect to the error rate as the number of hash functions increases.

**Fig. 11:**
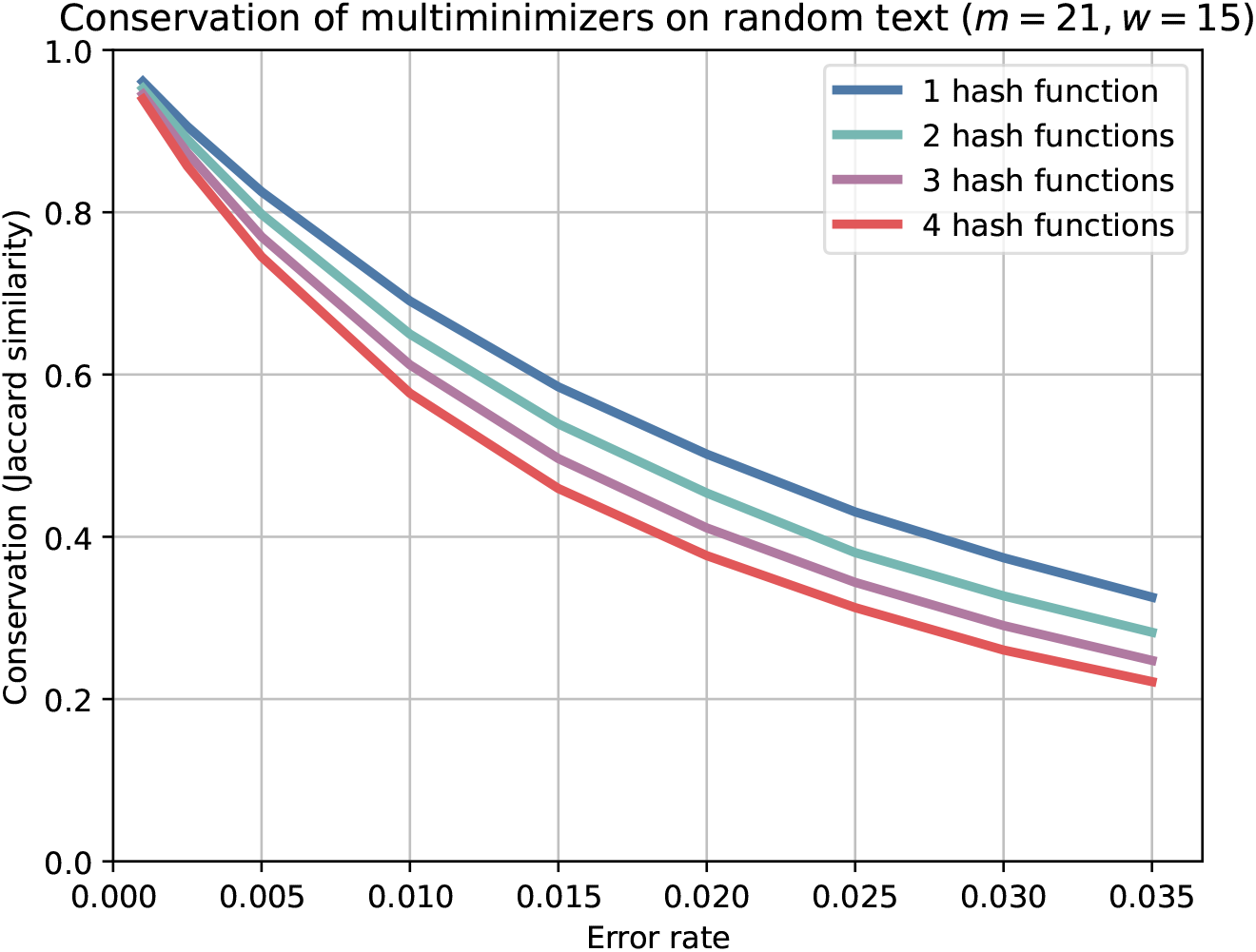
Conservation of sampled multiminimizers, measured by their Jaccard similarity, with 1 ≤ *N* ≤ 4 for a random string of size 10^5^ compared to a mutated version of the string with up to 3.5% of errors.

### S9 Linearity of indexation time

In this section, we show that the iteration over multiminimizers is linear with regard to the number of hash functions. The results are shown in Figure 12.

**Fig. 12:**
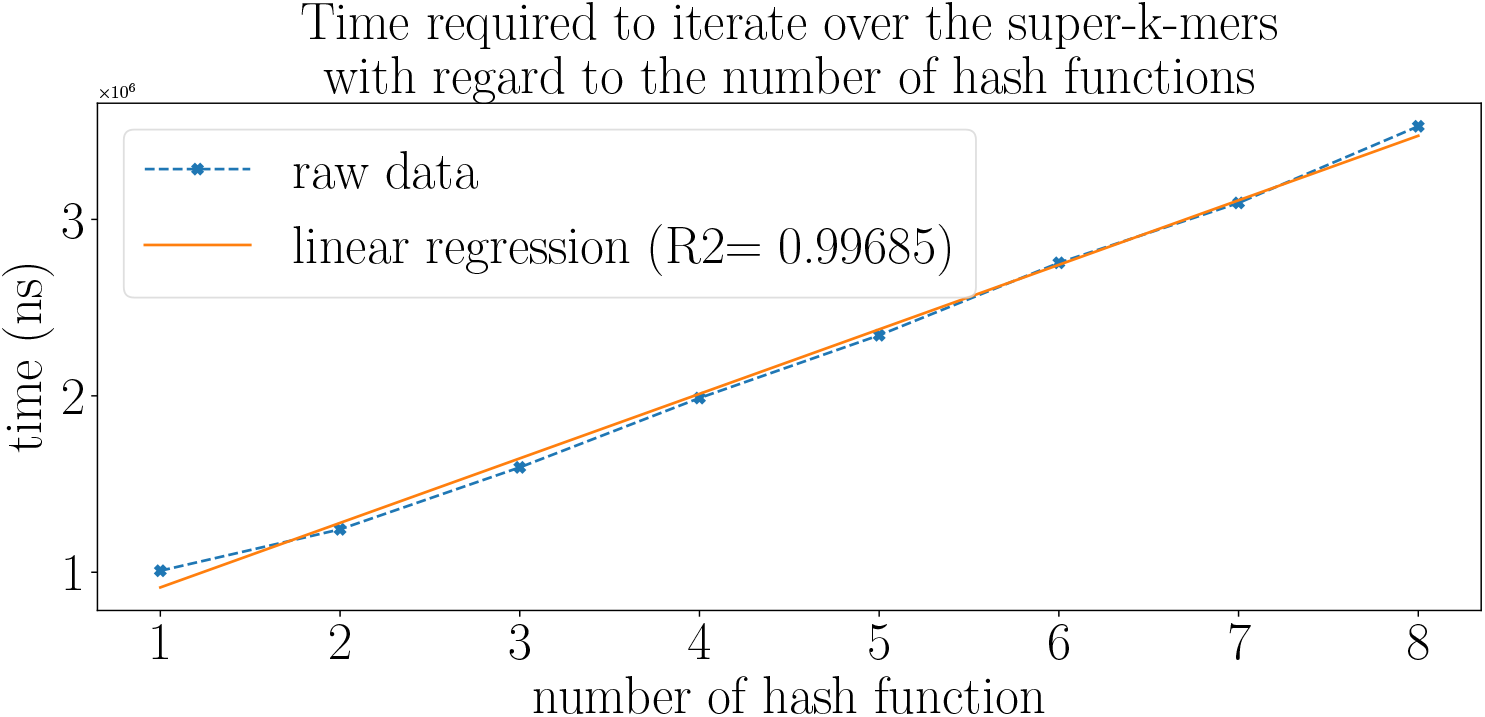
Time taken to index a random string of size 10^5^. The linear regression shows that the time taken to iterate over the multiminimizers is approximately the time needed to iterate over one random minimizer scheme times the number of hash functions.

## Footnotes

3 While the *positions* of minimizers are somehow uniformly distributed within a window, as seen in Section 2.2, it is not the case of theirs *values* — that are the ones that repeat and therefore lead to a decrease in *d*^∗^. Indeed, the values of the minimizers are distributed as min(*R*_1_, …, *R*_*w*_ ) where *R*_*i*_ are i.i.d. uniform variables, and therefore are strongly biased towards small values.

